# Boosting BDNF in muscle rescues impaired axonal transport in a mouse model of DI-CMTC peripheral neuropathy

**DOI:** 10.1101/2023.04.09.536152

**Authors:** Elena R. Rhymes, Rebecca L. Simkin, Ji Qu, David Villarroel-Campos, Sunaina Surana, Yao Tong, Ryan Shapiro, Robert W. Burgess, Xiang-Lei Yang, Giampietro Schiavo, James N. Sleigh

## Abstract

Charcot-Marie-Tooth disease (CMT) is a genetic peripheral neuropathy caused by mutations in many functionally diverse genes. The aminoacyl-tRNA synthetase (ARS) enzymes, which transfer amino acids to partner tRNAs for protein synthesis, represent the largest protein family genetically linked to CMT aetiology, suggesting pathomechanistic commonalities. Dominant intermediate CMT type C (DI-CMTC) is caused by *YARS1* mutations driving a toxic gain-of-function in the encoded tyrosyl-tRNA synthetase (TyrRS), which is mediated by exposure of consensus neomorphic surfaces through conformational changes of the mutant protein. In this study, we first showed that human DI-CMTC-causing TyrRS^E196K^ mis-interacts with the extracellular domain of the BDNF receptor TrkB, an aberrant association we have previously characterised for several mutant glycyl-tRNA synthetases linked to CMT type 2D (CMT2D). We then performed temporal neuromuscular assessments of *Yars*^*E196K*^ mice modelling DI-CMT. We determined that *Yars*^*E196K*^ homozygotes display a selective, age-dependent impairment in *in vivo* axonal transport of neurotrophin-containing signalling endosomes, phenocopying CMT2D mice. This impairment is replicated by injection of recombinant TyrRS^E196K^, but not TyrRS^WT^, into muscles of wild-type mice. Augmenting BDNF in DI-CMTC muscles, through injection of recombinant protein or muscle-specific gene therapy, resulted in complete axonal transport correction. Therefore, this work identifies a non-cell autonomous pathomechanism common to ARS-related neuropathies, and highlights the potential of boosting BDNF levels in muscles as a therapeutic strategy.

## Introduction

Charcot-Marie-Tooth disease (CMT) is a heritable, genetically diverse form of motor and sensory neuropathy that affects 1 in ≈2,500 people [58, 62]. Patients display slowly progressive weakness, muscle wasting and sensory dysfunction, which usually manifests in the hands and feet during adolescence, resulting in life-long disability and a significant health and financial burden on society [63]. CMT can be categorised into two main subtypes depending on pathogenesis and clinical assessment of nerve conduction velocity (NCV) in the median or ulnar nerve: demyelinating/type 1 CMT (CMT1) and axonal/type 2 CMT (CMT2). CMT1 is caused by the loss of myelinating Schwann cells that ensheath peripheral nerves leading to reduced NCV (<38 m/s), whereas CMT2 is caused by loss of peripheral axons, with little effect on NCV (>45 m/s) [57]. Some forms of genetic peripheral neuropathy display clinical features of CMT1 and CMT2, including intermediate NCVs (25-45 m/s), and are thus categorised as intermediate CMT [62], although this classification can vary [7].

Mutations in more than 100 genes are currently known to cause CMT, purely motor, or purely sensory inherited neuropathies [54]. These genes encode proteins involved in many cellular processes, including mitochondrial biogenesis, axonal transport, protein synthesis and endosomal sorting [60]. Many forms of CMT1 are caused by mutations in Schwann cell-critical proteins (e.g., peripheral myelin protein 22 and myelin protein zero), which provides a clear rationale for the selective neuropathology characteristic of CMT. However, the pathomechanisms driving motor and sensory dysfunction in CMT2 are less understood, in part, because the causative genes encode proteins required by many different cells and tissues. Indeed, the gene family linked to the most CMT subtypes encodes aminoacyl-tRNA synthetase (ARS) enzymes, which transfer specific amino acids to cognate tRNAs, and are therefore critical to protein synthesis [94].

Seven ARS genes have so far been linked to CMT (*AARS1, GARS1, HARS1, MARS1, SARS1, WARS1* and *YARS1*) with varying degrees of certainty about pathogenicity [1, 3, 33, 37, 38, 41, 88]. ARS-related CMT subtypes are all dominantly inherited and display a similar clinical phenotype. Although many ARS mutations affect tRNA aminoacylation, this is not a pre-requisite for disease [50, 94, 96]. Rather, a toxic gain-of-function appears to be the primary driver of neuropathology, at least in the well-studied *GARS1*-linked CMT type 2D (CMT2D) [29, 30, 44, 46, 47]. Indeed, point mutations in *GARS1* have been shown to cause a conformational opening of the encoded glycyl-tRNA synthetase (GlyRS), exposing consensus regions, which are normally buried in the wild-type enzyme, to the protein exterior [17, 34, 35]. These neomorphic surfaces enable aberrant protein-protein interactions with several intra- and extra-cellular binding partners, including neuropilin-1 [34], Trk receptors [68], HDAC6 [43] and G3BP1 [20]. Moreover, mutant GlyRS exhibits altered tRNA^Gly^ binding and release, hindering dissociation and causing protein synthesis defects linked to activation of the integrated stress response (ISR) [79, 97].

Neurotrophins are a family of secreted trophic factors that preferentially bind to neuronally-expressed Trk receptors (NGF to TrkA, BDNF and NT-4/5 to TrkB and NT-3 to TrkC), enabling local and long-range signalling critical to nerve cell homeostasis [16]. We have demonstrated that several different forms of human mutant GlyRS aberrantly associate with the extracellular domains (ECDs) of neurotrophin receptors, whereas GlyRS^WT^ does not [68, 76]. Further, we recently determined that this mis-interaction between mutant GlyRS and TrkB perturbs axonal transport of neurotrophin-containing signalling endosomes *in vivo*, and that treatment of CMT2D muscles with the TrkB ligand BDNF is able to fully rescue the transport disruption [76]. GlyRS is secreted from many different cell types, including neurons and muscle cells, and circulates freely in healthy mouse and human blood [28, 29, 34, 53], underpinning the significance of this mis-interaction *in vivo*.

Heterozygous point mutations in tyrosyl-tRNA synthetase (TyrRS)-encoding *YARS1* cause demyelinating and axonal neuropathy symptoms that result in a diagnosis of dominant intermediate CMT type C (DI-CMTC) [38]. Patients display mild to moderate disease severity with intermediate NCVs and predominant motor versus sensory involvement, which tends to be milder in females [83]. DI-CMTC is unlikely to result from haploinsufficiency, since a 50% reduction in *Yars* mRNA produces no overt phenotype in mice [36]. Additionally, children with severe, multi-system syndromes caused by biallelic *YARS1* loss-of-function mutations display no clear peripheral neuropathy [48, 87], arguing against a loss of *YARS1* function underlying DI-CMTC.

Similar to GlyRS, several different CMT mutations in *YARS1* result in an alternative stable conformation of TyrRS [10], which enhances its binding affinity for the transcriptional regulators TRIM28 and HDAC1 [9], as well as F-actin [23]. Moreover, a *Drosophila* model of DI-CMTC, in which mutant *YARS1* is overexpressed, displays a similar pattern of neuropathology and mRNA translation impairment in motor and sensory neurons to that caused by mutant *GARS1* [47, 81]. Furthermore, a new mouse model of DI-CMTC, which carries the E196K mutation in the catalytic domain of TyrRS (*Yars*^*E196K*^; see below), displays an activation of the ISR in α motor neurons similar to that identified in CMT2D mice, albeit to a lesser extent [79]. It therefore appears that *GARS1*- and *YARS1*-related neuropathies share key features, suggesting that common gain-of-function mechanism(s) may mediate neuropathology driven by mutations in the ARS-encoding genes.

To better understand DI-CMTC and provide the first mammalian model of the disease, the Burgess Laboratory engineered the Glu196Lys (c.586G>A; p.E196K) patient mutation into exon 5 of the mouse *Yars* gene (also c.586G>A), creating the *Yars*^*E196K*^ mouse [79]. The TyrRS^E196K^ mutation has little effect on tyrosine aminoacylation [25, 38, 81], hence *Yars*^*E196K*^ mice can be studied as heterozygotes or homozygotes, unlike mouse models for CMT2D [2, 44, 65]. On a C57BL/6N background, heterozygous *Yars*^*E196K/+*^ mice displayed little to no neuromuscular phenotype. However, homozygous *Yars*^*E196K/E196K*^ mice have impaired motor performance in the wire hang test that manifests at 2 months of age and marginally declines by 4 and 7 months [36]. Additionally, the *Yars*^*E196K/E196K*^ mutants exhibited impaired NCVs by 4 months, as well as reduced calibres of motor, but not sensory, axons without any axon loss or myelination defects up to 7 months. Whilst the homozygous mutant mice do not precisely model the human genetics of DI-CMTC, they provide an excellent opportunity to assess whether additional phenotypes are common between DI-CMTC and CMT2D mice, which would be expected if these forms of CMT share underlying pathomechanisms.

In this study, we therefore set out with two main aims: *1*) to determine whether the E196K mutation in *YARS1* triggers a pathological association with the ECD of TrkB similar to that found in CMT2D-linked *GARS1* mutants [68, 76]; and *2*) to provide further temporal characterisation of neuromuscular phenotypes in *Yars*^*E196K/+*^ and *Yars*^*E196K/E196K*^ mice to reveal neuropathological similarities between ARS-linked neuropathies in a mammalian setting.

## Materials and Methods

### Animals

Mice were maintained under a 12 h light/dark cycle at a constant room temperature of ≈21 °C with water and food *ad libitum* (Teklad global 18% protein rodent diet, Envigo, 2018C). Cages were enriched with nesting material, plastic/cardboard tubes and wooden chew sticks as standard, and occasionally with cardboard houses. *Yars*^*E196K*^ mice originally on a C57BL/6N background were backcrossed five times onto the C57BL/6J background and subsequently maintained as heterozygous × wild-type/heterozygous/homozygous breeding pairs. Both males and females were used, details of which are provided in **Table S1**. To reduce the number of animals used in this study, multiple assessments were performed on individual mice when possible; for instance, some mice were used for body weight and grip strength testing, prior to euthanising for dissection of muscles and sciatic nerves. Accordingly, a total of 182 mice was used across all experiments. Animals sacrificed for 3, 9 and 15 month time points were 86-99, 265-287 and 450-459 days old, respectively (see **Table S1**). 8 month-old *Yars*^*E196K/E196K*^ mice injected with adeno-associated viruses (AAVs) were aged 243-245 days. Post-natal day 1 (P1) was defined as the day after a litter was first found.

### Genotyping

DNA was extracted from ear clips as described previously [66] and the following primers were used in a single PCR for genotyping: Yars_F 5’-TGG CTC CAC CCT ATG AGA AC-3’, Yars^WT^_R 5’-AAA CCA CAC CAA CAG CCT TC-3’ and Yars^E196K^_R 5’-CCG CGT TAC TTC CTT TTC C-3’. PCR cycling conditions were as follows: 94 °C for 3 min, 35 × [94 °C for 30 s, 60 °C for 35 s, 72 °C for 1 min], 72 °C for 7 min. The wild-type and E196K sequences produce bands of 598 and 660 bp, respectively.

### *In vitro* pull-down assay

Co-immunoprecipitation experiments using the ECD of human TrkB-Fc (Cys32-His430; R&D Systems, 688-TK) and control human IgG-Fc (110-HG, R&D Systems) were performed in NSC-34 cells (CELLutions Biosystems, CLU140) transfected with plasmids encoding human TyrRS^WT^-V5, TyrRS^E196K^-V5 and GlyRS^ΔETAQ^-V5, as previously described [68, 69].

### Protein extraction and western blotting

Sciatic nerves and tibialis anterior muscles were dissected from PBS-perfused and non-perfused mice. Proteins were extracted from NSC-34 cells and tissues, and evaluated by western blotting as detailed previously [76]. Primary and secondary antibodies used for western blotting are listed in **Table S2** and **Table S3**, respectively. Densitometry was performed as previously described [66] using total protein stained with 0.1% Coomassie Brilliant Blue R-250 (Thermo Fisher, 20278) as the loading control [95]. Hook1 bands at ≈110 kDa and ≈90 kDa were quantified together in sciatic nerves analyses.

### Immunofluorescence

Dorsal root ganglia (DRG) from spinal levels lumbar 1 (L1) to L5 and cervical 4 (C4) to C8 were dissected [77], sectioned, stained and analysed as described elsewhere [68]. An average of 2,022 ± 96 lumbar cells (range 1,428-2,684) and 365 ± 23 cervical cells (range 279-526) across three to five DRG were assessed in each mouse. The percentages of NF200^+^ and peripherin^+^ cells per mouse were calculated by averaging values from individual DRG. Epitrochleoanconeous (ETA), flexor digitorum brevis (FDB) and hind/forelimb lumbrical muscles were dissected, processed for neuromuscular junction (NMJ) imaging, and analysed as detailed previously [67, 70, 91]. One hundred NMJs per mouse were analysed for assessment of maturation and degenerative phenotypes. Spinal cords were dissected from 4% paraformaldehyde-perfused mice and sectioned at 30 µm; choline acetyltransferase (ChAT)-positive neurons within the L3-L5 anterior horns were counted and their areas measured as previously described [76]. To assess eIF2α activation, spinal cords were processed in parallel; the intensity of phopho-eIF2α staining was determined relative to the intensity of ChAT staining for each motor neuron cell body, as performed elsewhere [79], before calculating an average per mouse. Primary and secondary antibodies used for immunofluorescence are listed in **Table S2** and **Table S3**, respectively. In addition, AlexaFluor555-conjugated α-bungarotoxin (α-BTX, Life Technologies, B35451) was used at 1:1,000 to identify post-synaptic acetylcholine receptors (AChRs). Fluorescent imaging of fixed samples and live nerves (see below) was performed on either an inverted LSM780 or LSM980 laser-scanning microscope (ZEISS).

### *In vivo* imaging of signalling endosome axonal transport

Live imaging of signalling endosome axonal transport was performed using an atoxic binding fragment of the tetanus neurotoxin (H_C_T, residues 875-1,315 fused to a cysteine-rich tag and a human influenza haemagglutinin epitope) purified and labelled as formerly described [27, 59]. *In vivo* imaging and analysis of axonal transport was performed as described previously [75, 76, 86], using a pre-warmed environmental chamber set to 38°C and acquiring images every ≈3 seconds using a 63× Plan-Apochromat oil immersion objective (ZEISS). For phenotyping at 3, 9 and 15 months, H_C_T was injected under isoflurane-induced anaesthesia into the tibialis anterior on one side of the body and the contralateral gastrocnemius; this enabled transport assessment in peripheral nerves supplying distinct hindlimb muscles of the same mouse. To assess the impact of TyrRS on transport, 25 ng/muscle of recombinant human proteins (TyrRS^WT^ or TyrRS^E196K^, produced as described previously [23]) were pre-mixed with H_C_T prior to bilateral administration into the tibialis anterior muscles of wild-type mice. The side of injection (tibialis anterior versus gastrocnemius, and TyrRS^WT^ versus TyrRS^E196K^) was alternated between mice to eliminate time-under-anaesthesia and right/left biases. Imaging was performed 4-8 h later in both sciatic nerves. By selecting thicker H_C_T-positive axons, evaluation of axonal transport in motor axons was prioritised [74]. To assess trafficking dynamics, 15 endosomes per axon and three to five axons per mouse were manually tracked using the TrackMate plugin on ImageJ (http://rsb.info.nih.gov/ij/) [84]. Endosome reversals were infrequent (<1% of all frame-to-frame movements) and recorded as positive values. Endosome frame-to-frame speeds are presented in frequency histograms; a mean of 393 ± 7 movements (range 279-716) per animal were assessed (in a total of 130 mice across the study).

### Body weight and grip strength testing

Animals were weighed, and all-limb grip strength was assessed using a Grip Strength Meter (Bioseb, GS3). Mice were assessed on two separate days, on which three trials (separated by at least 5 min) were performed. The maximum recorded values from each of the two days were used to calculate a mean grip strength per animal. Relative body weight and grip strength were determined by generating the mean value for wild-type mice of each sex and then calculating individual mouse values as a percentage of the wild-type mean of their respective sex.

### AAV8-tMCK virus production

Self-complementary AAV (scAAV) expression plasmids were created by OXGENE (Oxford, UK); the incorporated sequences of the 745 bp tMCK promoter, 714 bp eGFP and 744 bp human pre-proBDNF are provided elsewhere [76]. pSF-scAAV-tMCK-eGFP and pSF-scAAV-tMCK-BDNF plasmids were packaged into AAV serotype 8 particles by Charles River Laboratories (formerly Vigene Biosciences, Rockville, MD, USA). Viruses were stored at -80 °C in 0.01% (v/v) pluronic F68 surfactant in PBS and diluted in sterile PBS for injection.

### Intramuscular injections of BDNF and AAV

To assess the impact of boosting muscle BDNF on endosome axonal transport, bilateral injections into the tibialis anterior muscles of *Yars*^*E196K/E196K*^ mice were performed under isoflurane-induced anaesthesia. Acute treatment was evaluated in 9 month-old mice by pre-mixing and co-administering 5 µg H_C_T-555 with 25 ng recombinant human BDNF (Peprotech, 450-02) into one tibialis anterior muscle and 5 µg H_C_T-555 with PBS vehicle into the contralateral tibialis anterior, both in a volume of ≈2.5 µl per muscle. For analysing longer term treatment, 10 µl bilateral injections of 2.0 × 10^11^ vg AAV8-tMCK-eGFP into one tibialis anterior and 2.0 × 10^11^ vg AAV8-tMCK-BDNF into the contralateral tibialis anterior were performed in 8 month-old mice. 29-30 days post-AAV treatment, *in vivo* axonal transport of signalling endosomes was assessed at 9 months as described above. For these treatments, the side of injection was alternated between mice to avoid biases.

### Statistics

Data were assumed to be normally distributed unless evidence to the contrary were provided by the Kolmogorov-Smirnov test for normality, while equal variance between groups was assumed. The Bonferroni correction was applied to the Kolmogorov-Smirnov tests within each experiment. Normally-distributed data were analysed using unpaired and paired *t*-tests or one-way and two-way analysis of variance (ANOVA) tests followed by Šídák’s multiple comparisons test. Non-normally distributed data were analysed using a Mann-Whitney *U* test, Wilcoxon matched-pairs signed rank test or Kruskal-Wallis test followed by Dunn’s multiple comparisons test. Sample sizes, which were pre-determined using power calculations and previous experience [67, 68, 70-72, 74, 76], are reported in figure legends and represent biological replicates (*i*.*e*., individual animals). Means ± SEM are plotted for all graphs. All tests were two-sided and an α-level of *P* < 0.05 was used to determine significance. GraphPad Prism 10 software (version 10.1.1) was used for statistical analyses and figure production.

## Results

### TyrRS^E196K^ mis-interacts with the extracellular domain of TrkB

Our initial aim in this study was to test whether the TyrRS^E196K^ mutant aberrantly interacts with the ECD of TrkB. To do this, we transfected NSC-34 cells with constructs encoding V5-tagged human TyrRS^WT^, TyrRS^E196K^ or GlyRS^ΔETAQ^, with the latter included as a positive control [76]. Cells were subsequently lysed and co-immunoprecipitation experiments performed with the ECD of Fc-tagged TrkB. Western blotting of pull-downs performed in triplicate revealed that TyrRS^E196K^ does indeed mis-associate with TrkB to a higher extent than TyrRS^WT^, but less than GlyRS^ΔETAQ^ (**Figure 1A-B**). Importantly for a potential shared pathomechanism with GlyRS, TyrRS is also secreted and is found in blood [13, 31, 64].

**Figure 1.**
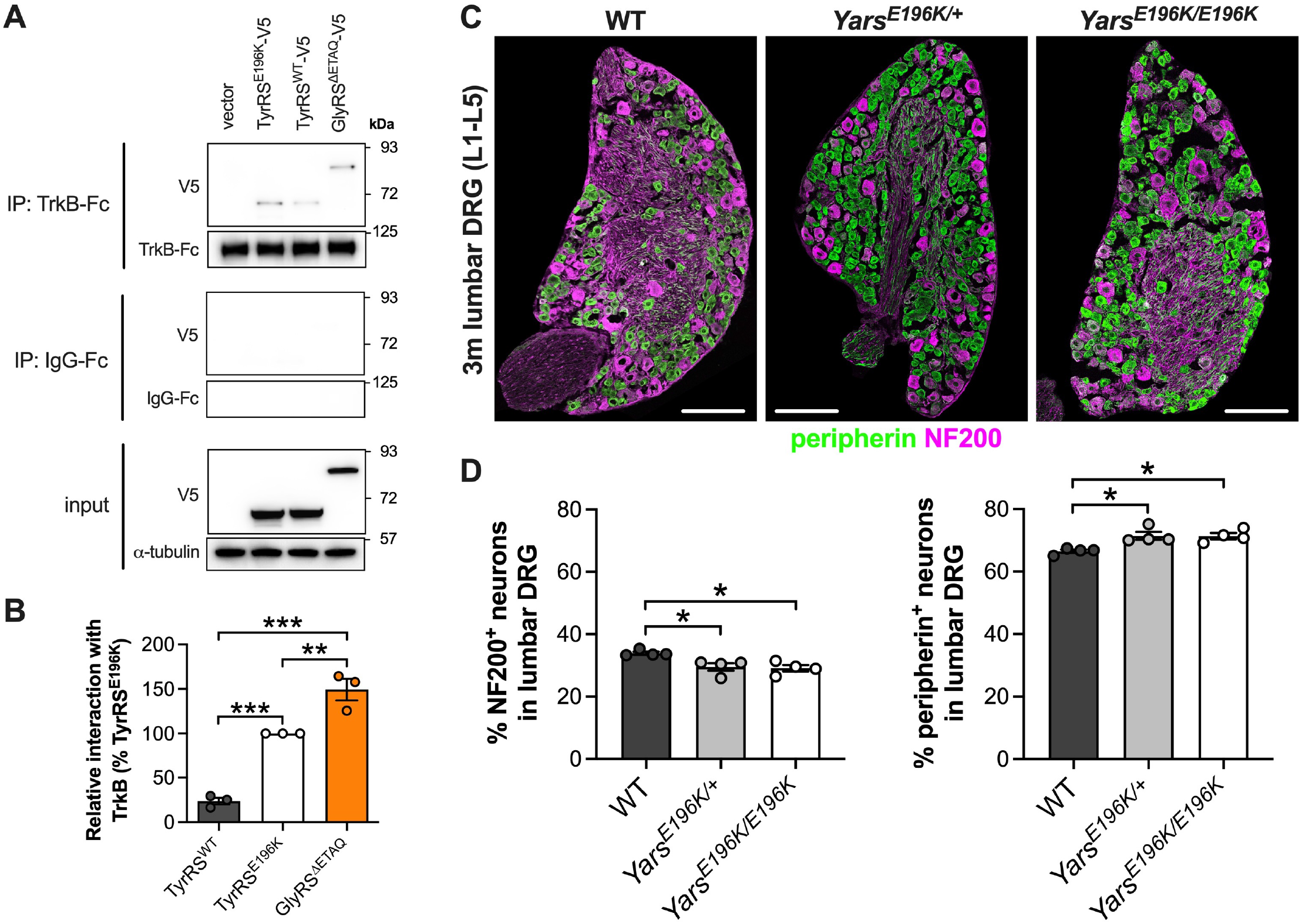
TyrRS^E196K^ aberrantly interacts with the extracellular domain of TrkB underlying altered sensory neuron subtype proportions. (**A**) Representative western blots of *in vitro* pull-downs from NSC-34 cells showing that human TyrRS^E196K^ interacts with the extracellular domain of human TrkB-Fc to a greater extent than TyrRS^WT^. GlyRS^ΔETAQ^ was used as a positive control. *IP*, immunoprecipitation; *kDa*, kilodalton. (**B**) Quantification of the relative interactions between TyrRS^WT^, TyrRS^E196K^ and GlyRS^ΔETAQ^ with TrkB-Fc (*P* < 0.001 one-way ANOVA). *n* = 3. (**C**) Representative immunofluorescent images of 3 month-old wild-type, *Yars*^*E196K/+*^ and *Yars*^*E196K/E196K*^ lumbar DRG sections (10 µm) stained for NF200 (magenta) and peripherin (green). Scale bars = 100 µm. (**D**) *Yars*^*E196K/+*^ and *Yars*^*E196K/E196K*^ mice display a lower percentage of NF200^+^ neurons (left graph, *P* = 0.013 Kruskal-Wallis test) and a higher percentage of peripherin^+^ neurons (right graph, *P* = 0.013 Kruskal-Wallis test) in lumbar 1 (L1) to L5 DRG compared to wild-type. *n* = 4. For all graphs, \**P* < 0.05, \*\**P* < 0.01, \*\*\**P* < 0.001 Šidák’s/Dunn’s multiple comparisons test; means ± standard error of the mean (SEM) plotted. See also **Figure S1**.

### Yars^E196K^ *mice display a perturbation in sensory neuron identity*

We have previously revealed that lumbar, but not cervical DRG of CMT2D mice display fewer NF200^+^ mechanoreceptors/proprioceptors, and a concomitant increase in the percentage of peripherin^+^ nociceptors [68, 71]. During pre-natal development, sensory neurons can express multiple Trk receptors, reaching their mature Trk identity and function after birth [40]. The neuropathic DRG phenotype is thus perhaps linked to the aberrant interaction between mutant GlyRS and the Trk receptors causing a subversion of sensory neuron differentiation.

Given that mutant TyrRS mis-associates with TrkB, we wanted to determine whether *Yars*^*E196K*^ mice display a similar alteration in sensory neuron populations. We therefore dissected lumbar DRG from 3 month-old wild-type, *Yars*^*E196K/+*^ and *Yars*^*E196K/E196K*^ mice and performed immunohistochemical analysis of NF200 and peripherin (**Figure 1C**). Compared to wild-type littermate controls, we identified that both *Yars*^*E196K/+*^ and *Yars*^*E196K/E196K*^ mice display a lower percentage of NF200^+^ neurons and an increase in the percentage of peripherin^+^ neurons (**Figure 1D**). Similar to mutant *Gars* mice, this phenotype was not present in *Yars*^*E196K/+*^ cervical DRG (**Figure S1**). Heterozygous and homozygous mutant *Yars* mice thus both phenocopy CMT2D mice in the spinal region-specific perturbation of sensory neuron identity.

To evaluate the severity of this sensory phenotype compared to the change observed in CMT2D models, the percentage of NF200^+^ neurons in lumbar DRG relative to wild-type (*n*.*b*., not raw percentages) was calculated for *Yars*^*E196K/+*^ (87.1 ± 3.5%) and *Yars*^*E196K/E196K*^ (85.8 ± 3.0%), as well as 3 month-old *Gars*^*ΔETAQ/+*^ mice (74.7 ± 1.3%; data not shown). All DRG assessments were performed by the same researcher (RLS), reducing variability, but relative values were required because the experiments were not performed or analysed in parallel. Statistical comparisons between the three ARS CMT genotypes indicate a difference (\**P* = 0.020 Kruskal-Wallis test), and that the reduction in NF200^+^ cells is greater in *Gars*^*ΔETAQ/+*^ lumbar DRG than *Yars*^*E196K/E196K*^ (\**P* = 0.025 Dunn’s multiple comparisons test). Therefore, the severity of disruption in sensory neuron identity correlates with the extent of the mis-interaction between the mutant ARS protein and TrkB.

### *Body weight, NMJ innervation and grip strength are all unaffected in* Yars^E196K^ *mice at 3 months*

The above DRG analyses, and all subsequent experiments, were performed in *Yars*^*E196K*^ mice on a C57BL/6J background, which was chosen to allow for direct comparison with the CMT2D strains that are maintained on the same background in our laboratory. In the original assessment, when *Yars*^*E196K*^ mice were generated, experiments were performed at 2, 4 and 7 months of age and the mutation was maintained on a slightly different genetic background (C57BL/6N) [36]. In the study presented here, we chose to evaluate *Yars*^*E196K*^ mice at 3 and 9 months to extend the phenotyping age range and allow for evaluation of similarities with mutant *Gars* alleles, which we have extensively characterised at 3 months [68, 70, 72, 76].

First, we assessed the body weight of 3 month-old wild-type, *Yars*^*E196K/+*^ and *Yars*^*E196K/E196K*^ (**Figure S2**). We observed no differences in body weight or relative body weight of females or males, corroborating previous analyses at 4 and 7 months [36]. At 4 months, *Yars*^*E196K/E196K*^ mice have reduced NCVs, thinner motor axons and impaired motor function and endurance, as assessed by the wire hang (inverted grid) test [36]. We therefore analysed the architecture of NMJs and the maximum grip strength of *Yars*^*E196K*^ mice. We have previously identified a spectrum of vulnerability to degeneration of NMJs in different muscles in *Gars*^*C201R/+*^ mice modelling CMT2D [70, 72]. At 3 months of age, the ETA muscle in the forearm is almost completely unaffected (≈4% denervation); in contrast, the hindlimb lumbricals and FDB muscles of the hindpaw display ≈43% and ≈63% denervated NMJs, respectively, whilst the forelimb lumbricals have an intermediate phenotype with ≈21% denervation. A similar pattern of vulnerability is also observed in *Gars*^*ΔETAQ/+*^ mice at 3 months (data not shown).

These same four muscles were dissected from 3 month-old *Yars*^*E196K*^ mice and stained to assess NMJ architecture using combined antibodies against SV2/2H3 to identify motor neurons, and fluorescently labelled α-BTX to label post-synaptic AChRs (**Figure S3A**). The extent of distal motor neuron pathology can be inferred by the degree of overlap between the pre- and post-synaptic staining and categorising NMJs as either fully innervated, partially denervated or vacant. We observed no NMJ denervation at 3 months of age in *Yars*^*E196K/+*^ or *Yars*^*E196K/E196K*^ mice in any of the four muscles (**Figure S3B**). Moreover, the delay in NMJ maturation present in CMT2D mice, assessed by calculating the percentage of NMJs with multiple innervating motor neurons (*i*.*e*., polyinnervation) [70, 72], was also largely absent in *Yars*^*E196K*^ muscles (**Figure S3C**). That is, except for a small, but significant, difference between forelimb lumbricals of heterozygous and homozygous mutants, which is likely a product of the lack of variability in the *Yars*^*E196K/E196K*^ mouse sample, rather than being biologically meaningful.

Consistent with the lack of morphological disruption at the NMJ, we were unable to detect any difference in grip strength between wild-type and *Yars*^*E196K*^ genotypes in females or males (**Figure S4**). This seemingly contrasts with previously performed wire hang assessments at 2 and 4 months in *Yars*^*E196K/E196K*^ mice [36]; however, the wire hang test measures muscle endurance whereas grip strength provides a readout of maximum strength.

### Yars^E196K/E196K^ *motor neurons display reduced areas and increased ISR activation at 3 months*

Motor neurons in the lumbar spinal cord of CMT2D mice are preserved, but display a reduction in area [76]. To assess whether *Yars*^*E196K*^ mice have a similar phenotype, the L3-L5 spinal cord was sectioned and stained for ChAT to identify motor neurons in the anterior horns and p-eIF2α to assess ISR activation (**Figure 2A**). We found that motor neuron numbers are similar between genotypes (**Figure 2B**), while *Yars*^*E196K*^ homozygotes displayed a reduction in motor neuron area (**Figure 2C**), similar to *Gars*^*C201R/+*^ mice. As body weight is unaffected at this age, the reduction in cell area is independent from body size and is concordant with the smaller motor axon calibres identified in *Yars*^*E196K/E196K*^ at 4 months [36]. We also confirmed a clear increase in p-eIF2α staining within motor neurons of *Yars*^*E196K/E196K*^ (**Figure 2D**), replicating in the C57BL/6J background previous results obtained in this neuropathy model [36].

**Figure 2:**
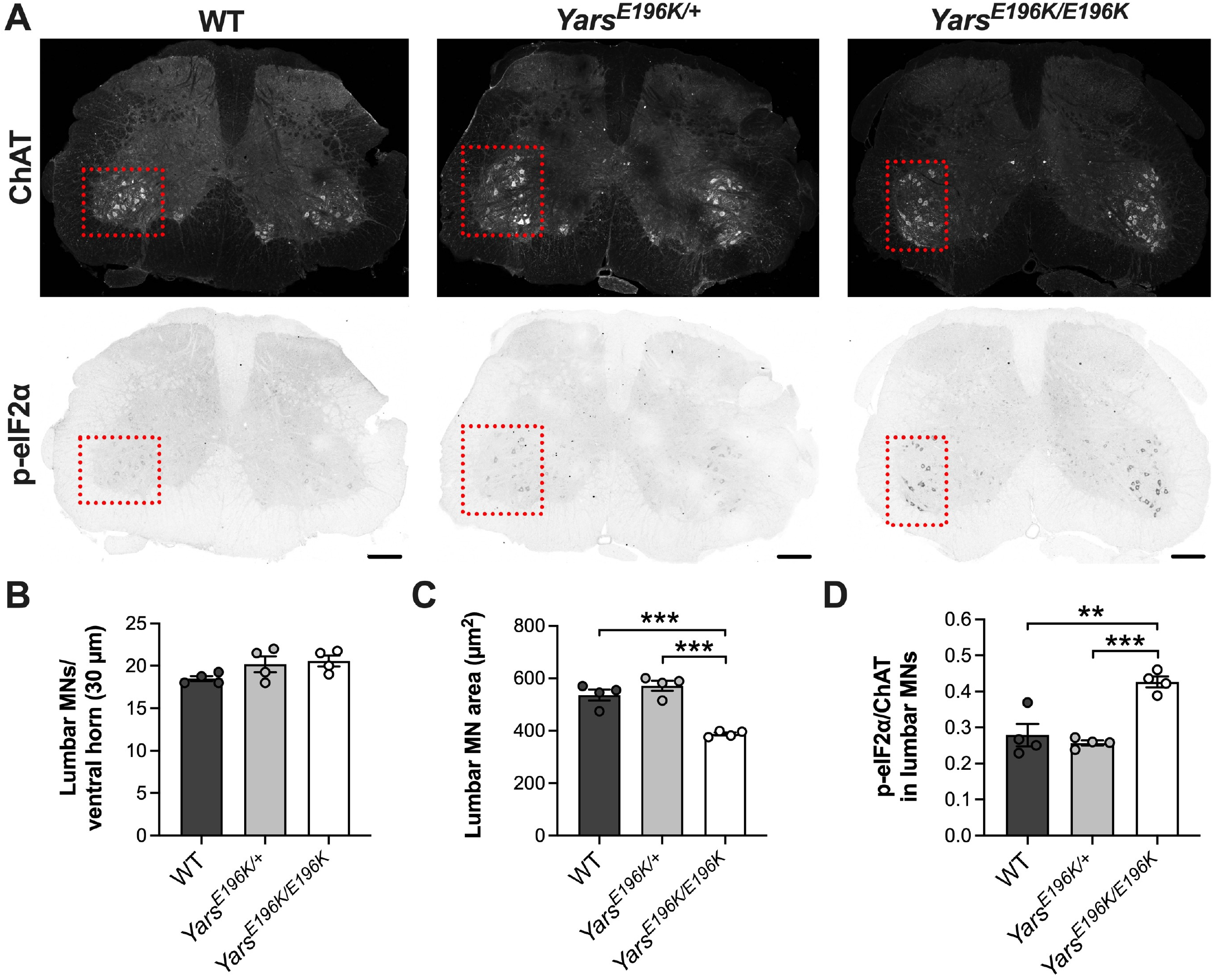
Motor neurons of homozygous *Yars*^*E196K*^ mice have reduced area at 3 months. (**A**) Representative immunofluorescent images of 3 month-old wild-type, *Yars*^*E196K/+*^ and *Yars*^*E196K/E196K*^ lumbar spinal cord sections (30 µm) stained for ChAT (top) and p-eIF2α (bottom). The red dashed line boxes highlight anterior horns containing ChAT^+^ motor neurons. Scale bars = 200 µm. (**B**) *Yars*^*E196K*^ mice display no lumbar motor neuron (MN) loss (*P* = 0.130 one-way ANOVA). (**C-D**) Motor neurons from homozygous, but not heterozygous *Yars*^*E196K*^ mice display a smaller area than wild-type (C, *P* < 0.001 one-way ANOVA) and have increased levels of p-eIF2α (D, *P* < 0.001 one-way ANOVA), indicative of activation of the integrated stress response (ISR). For all graphs, *n* = 4; \*\**P* < 0.01, \*\*\**P* < 0.001 Šidák’s multiple comparisons test; means ± SEM plotted.

### In vivo *axonal transport of signalling endosomes is unperturbed in* Yars^E196K^ *mice at 3 months*

*Gars*^*C201R/+*^ mice display impaired *in vivo* axonal transport of neurotrophin-containing signalling endosomes at 1 and 3 months of age, which manifests as a reduction in endosome trafficking speed after P15-16 [76]. This defect was identified by injecting a fluorescent retrograde probe (H_C_T) into the tibialis anterior and gastrocnemius muscles of anaesthetised mice, followed by time-lapse confocal imaging of the exposed sciatic nerve under terminal anaesthesia [75]. H_C_T is internalised at the NMJ and then sorted into signalling endosomes for retrograde axonal transport within motor axons [8]. Individual endosomes can be imaged, tracked and their dynamic properties assessed.

We therefore used H_C_T to identify whether *Yars*^*E196K*^ mice display an impairment in endosome axonal transport at 3 months of age (**Figure 3A**). However, rather than injecting both the tibialis anterior and gastrocnemius muscles together, we performed assessments of transport separately in these two muscles, as we have recently determined that their innervating axons can be differentially affected by disease in a mouse model of amyotrophic lateral sclerosis [85]. Importantly, the most common clinical signs reported in a small cohort of familial DI-CMTC patients were weakness and atrophy of the tibialis anterior and gastrocnemius [52], thus these two muscles are clinically relevant in the disease. However, in motor neurons innervating the tibialis anterior muscle, we observed no differences between genotypes in signalling endosome speed (**Figure 3B-C**), the percentage of time that endosomes remained stationary (**Figure 3D**) or the percentage of endosomes that paused (**Figure 3E**). A similar pattern was observed in gastrocnemius-innervating motor neurons (**Figure 3F-I**). Together, these results indicate that at 3 months of age, *Yars*^*E196K*^ mice display no disruption in axonal transport of signalling endosomes, commensurate with the body weight, NMJ and grip strength analyses.

**Figure 3.**
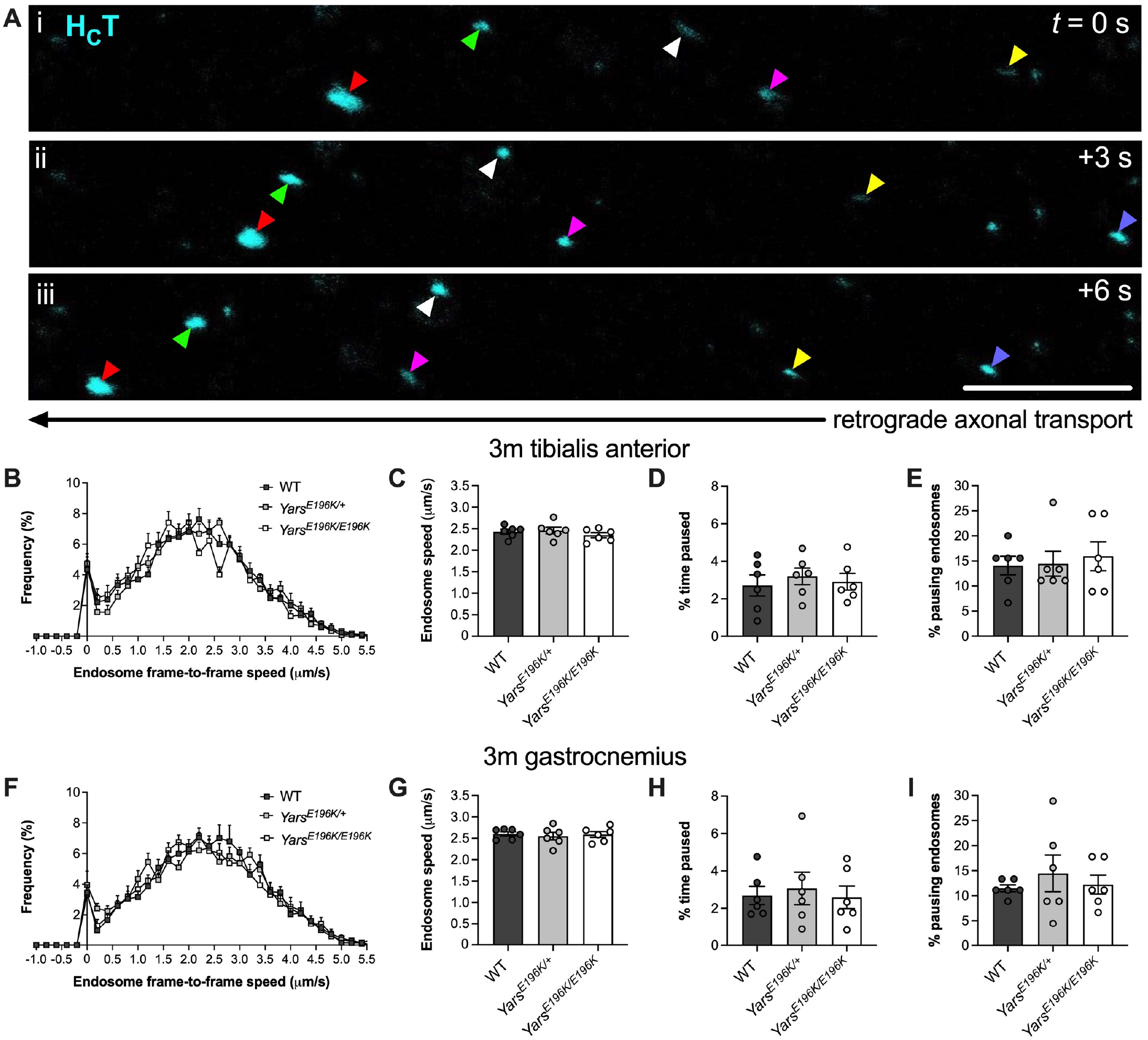
*In vivo* axonal transport of signalling endosomes is unaffected in *Yars*^*E196K*^ mice at 3 months. (**A**) Representative time-lapse confocal images of signalling endosomes labelled with an atoxic fluorescent fragment of tetanus neurotoxin (H_C_T, cyan) being retrogradely transported within peripheral nerve axons of live, anaesthetised mice. H_C_T-positive endosomes are individually tracked to quantitatively assess their dynamics. Colour-coded arrowheads identify six different endosomes. Scale bar = 10 µm. (**B**) Frame-to-frame speed histogram of signalling endosomes being transported within motor neurons innervating the tibialis anterior muscle of wild-type, *Yars*^*E196K/+*^ and *Yars*^*E196K/E196K*^ mice aged 3 months. (**C-E**) There is no difference between genotypes in signalling endosome speed (C, *P* = 0.487), the percentage of time paused (D, *P* = 0.781) or the percentage of pausing endosomes (E, *P* = 0.919 Kruskal-Wallis test) in motor neurons innervating the tibialis anterior muscle. (**F**) Frame-to-frame speed histogram of signalling endosomes being transported within motor neurons innervating the gastrocnemius muscle of wild-type, *Yars*^*E196K/+*^ and *Yars*^*E196K/E196K*^ mice aged 3 months. (**G-I**) There is no difference between genotypes in signalling endosome speed (G, *P* = 0.871), the percentage of time paused (H, *P* = 0.873) or the percentage of pausing endosomes (I, *P* = 0.671) in motor neurons innervating the gastrocnemius muscle. Endosomes within tibialis anterior- and gastrocnemius-innervating axons were analysed in the same mice. For all graphs, genotypes were compared using one-way ANOVAs, unless otherwise stated; *n* = 6; means ± SEM plotted. See also **Figure S8**.

We have previously determined that signalling endosome trafficking impairment in CMT2D mice correlates with reduced availability in the sciatic nerve of motor adaptor proteins critical to endosome transport [76]. As we see no transport deficits in the sciatic nerve of *Yars*^*E196K*^ mice at 3 months, we hypothesised that there would be no corresponding change in the levels of the adaptor proteins Snapin [12], Hook1 [49] or RILP [39], which have been shown to play a role in signalling endosome transport. We also assessed the levels of dynein intermediate chain (DIC), a major component of the retrograde motor protein cytoplasmic dynein [56], and probed for dynein intermediate chain phosphorylation at residue S81 (p-DIC), which facilitates the selective recruitment of the motor to signalling endosomes for retrograde trafficking upon Trk activation [42]. Contrary to expectation, we saw that DIC phosphorylation was reduced in *Yars*^*E196K/+*^ sciatic nerves, alongside an increase in RILP levels. In *Yars*^*E196K/E196K*^ nerves, dimeric Snapin (dSnapin) levels were reduced, whereas p-DIC, Hook1 and RILP were all increased (**Figure 4**). Although the cause of these changes is unknown, it is enticing to speculate that there may be compensatory mechanisms in endosome adaptor availability that ensure signalling endosome transport remains unaffected in mutant animals at 3 months of age.

**Figure 4.**
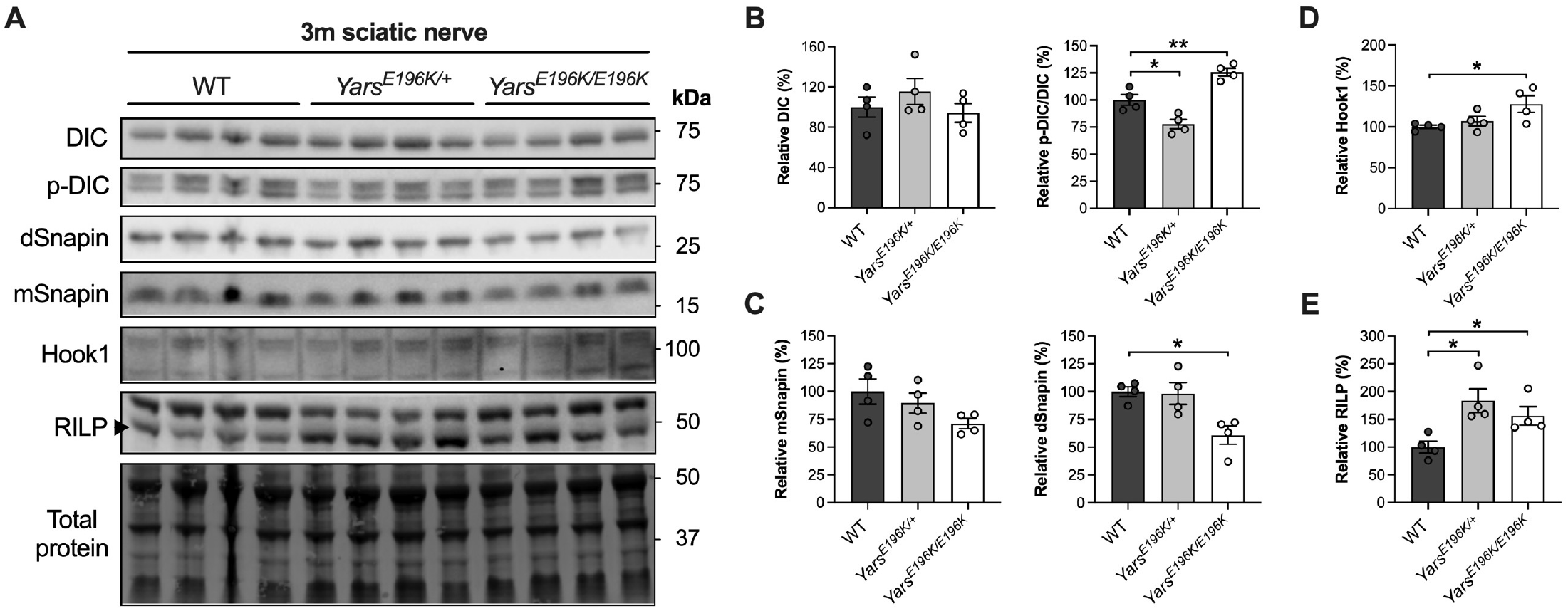
Complementary changes in endosome adaptor levels are observed in *Yars*^*E196K*^ sciatic nerves at 3 months. (**A**) Western blots of dynein intermediate chain (DIC), phosphorylated DIC (p-DIC) and dynein adaptor proteins Snapin, Hook1 and RILP in sciatic nerves from 3 month-old wild-type, *Yars*^*E196K/+*^ and *Yars*^*E196K/E196K*^ mice. *kDa*, kilodalton. (**B-E**) Densitometric analyses of DIC (B, left, *P* = 0.399), p-DIC relative to DIC (B, right, *P* < 0.001), monomeric Snapin (mSnapin, C, left, *P* = 0.113), dimeric Snapin (dSnapin, C, right, *P* < 0.01), Hook1 (D, *P* = 0.046) and RILP (E, *P* = 0.018). The ratio of p-DIC to DIC is reduced in *Yars*^*E196K/+*^ and increased in *Yars*^*E196K/E196K*^, the level of dSnapin is reduced and Hook1 increased in *Yars*^*E196K/E196K*^, and RILP is increased in both *Yars*^*E196K*^ genotypes. For all graphs, genotypes were compared using one-way ANOVAs; *n* = 4; \**P* < 0.05, \*\**P* < 0.01 Šídák’s multiple comparisons test; means ± SEM plotted.

Summarising the phenotype of *Yars*^*E196K*^ mice on a C57BL/6J background at 3 months, we observed a perturbation in sensory neuron identity, reduced lumbar motor neuron cell body areas and changes in adaptor protein levels in the sciatic nerve, but no deficits in body weight, NMJ innervation, maximal grip strength or *in vivo* axonal transport of signalling endosomes.

### Yars^E196K/E196K^ *mice display marginally reduced body weight, but normal motor innervation and function at 9 months*

As we detected only subtle changes in *Yars*^*E196K*^ mice at 3 months of age, we decided to re-assess mice at 9 months of age. We observed a small reduction in body weight and relative body weight of 9 month-old *Yars*^*E196K/E196K*^ females compared to *Yars*^*E196K/+*^, but not wild-type mice of the same sex (**Figure S5A-B**). Males were unaffected, suggesting that there may be a subtle sex-specific defect. When relative body weights of females and males were combined, we also saw a mild deficit in *Yars*^*E196K/E196K*^ compared to *Yars*^*E196K/+*^, but again not in comparison to wild-type animals (**Figure S5C**).

Mirroring the lack of a large reduction in body weight, we observed no degeneration of NMJs across the same four muscles analysed at 3 months (**Figure S6**). Of the six *Yars*^*E196K/E196K*^ mice analysed, five were females, indicating that there is unlikely to be sex-specific denervation. This is supported by the finding that *Yars*^*E196K*^ females and males display normal grip strength at 9 months (**Figure S7**). Again replicating the scenario at 3 months, we detected no lumbar motor neuron loss at 9 months and a similar reduction in cell body areas (**Figure 5**).

**Figure 5.**
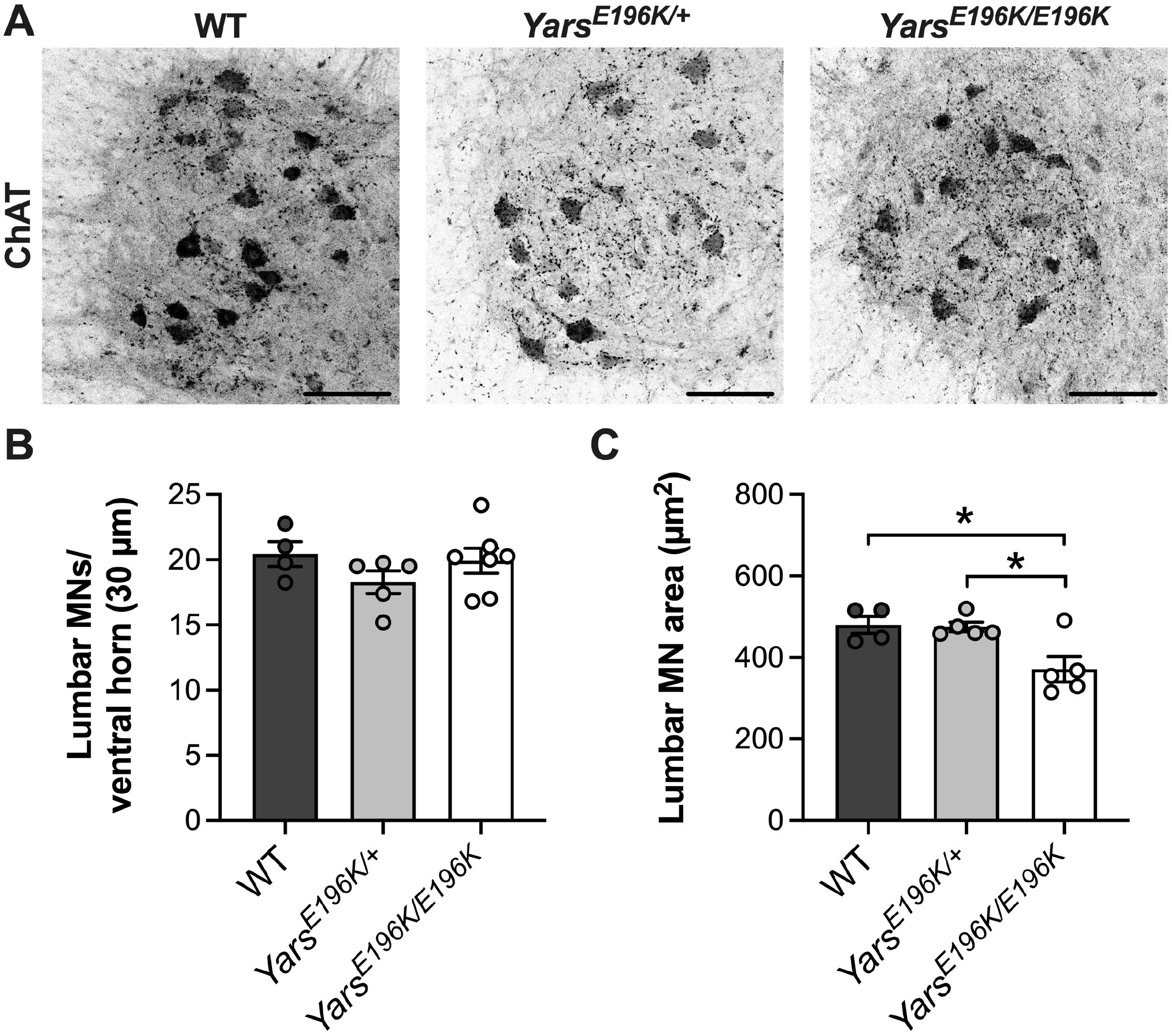
Motor neuron cell body areas are reduced at 9 months in *Yars*^*E196K*^ homozygotes. (**A**) Representative immunofluorescent images of 9 month-old wild-type, *Yars*^*E196K/+*^ and *Yars*^*E196K/E196K*^ lumbar spinal cord sections (30 µm) stained for ChAT. Scale bars = 100 µm. (**B**) *Yars*^*E196K*^ mice display no lumbar motor neuron (MN) loss (*P* = 0.320 one-way ANOVA). *n* = 4-7. (**C**) Motor neurons from homozygous *Yars*^*E196K*^ mice exhibit smaller cell body areas than *Yars*^*E196K/+*^ and wild-type mice (C, *P* = 0.010 one-way ANOVA). \**P* < 0.05 Šidák’s multiple comparisons test. *n* = 4-5. For both graphs, means ± SEM plotted.

### *In vivo* endosome *axonal transport is defective in* Yars^E196K/E196K^ *mice at 9 months*

We next evaluated *in vivo* axonal transport of signalling endosomes in motor axons innervating the tibialis anterior and gastrocnemius muscles of 9 month-old mice. *Yars*^*E196K/+*^ mice were indistinguishable from wild-type (**Figure 6** and **Figure S8**). However, *Yars*^*E196K/E196K*^ mice displayed a reduction in endosome trafficking speeds in tibialis anterior-innervating neurons, coupled with an increase in the percentage of pausing endosomes (**Figure 6A-D** and **Figure S8A-B**). This deficit appears to be selective to neurons innervating the tibialis anterior, as we saw no such impairments in gastrocnemius-innervating neurons (**Figure 6E-H** and **Figure S8C**).

**Figure 6.**
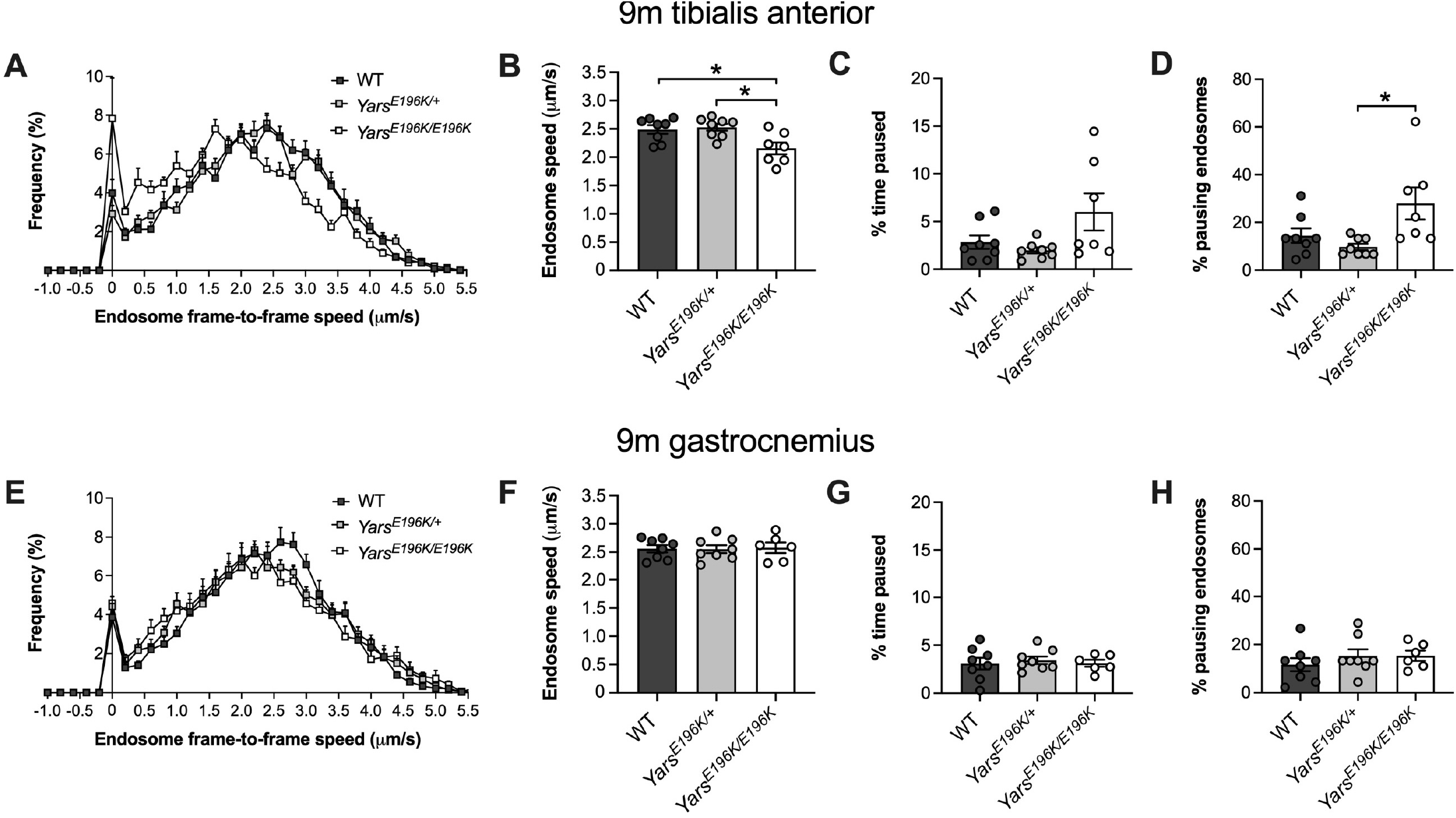
*In vivo* axonal transport is selectively impaired in *Yars*^*E196K*^ homozygotes at 9 months. (**A**) Frame-to-frame speed histogram of signalling endosomes being transported within motor neurons innervating the tibialis anterior muscle of wild-type, *Yars*^*E196K/+*^ and *Yars*^*E196K/E196K*^ mice aged 9 months. (**B**) Signalling endosome speed is reduced in tibialis anterior-innervating axons of *Yars*^*E196K/E196K*^ mice (*P* = 0.007). (**C-D**) The percentage of time paused is unaffected (C, *P* = 0.053), but the percentage of pausing endosomes is increased (D, *P* = 0.014) in tibialis anterior-innervating axons of *Yars*^*E196K/E196K*^ mice. (**E**) Frame-to-frame speed histogram of signalling endosomes being transported within motor neurons innervating the gastrocnemius muscle of wild-type, *Yars*^*E196K/+*^ and *Yars*^*E196K/E196K*^ mice aged 9 months. (**F-H**) There is no difference between genotypes in signalling endosome speed (F, *P* = 0.971), percentage of time paused (G, *P* = 0.848) or the percentage of pausing endosomes (H, *P* = 0.529 Kruskal-Wallis test) in motor neurons innervating the gastrocnemius. Endosomes within tibialis anterior- and gastrocnemius-innervating axons were analysed from the same mice. For all graphs, genotypes were compared using one-way ANOVAs, unless otherwise stated; *n* = 6-8; \**P* < 0.05 Šidák’s multiple comparisons test; means ± SEM plotted. See also **Figure S8** and **Figure S9**.

To assess whether the heterozygous mutants develop transport dysfunction with increased age, we assessed endosome dynamics at 15 months in both tibialis anterior- and gastrocnemius-innervating motor neurons in *Yars*^*E196K/+*^ mice (**Figure S9**). There was a small, but significant increase in the percentage of time paused at 15 months compared with 9 months; however, there was no reduction in endosome speed or the percentage of pausing cargoes, indicating that axonal transport of signalling endosomes remains largely unaffected in aged *Yars*^*E196K/+*^ mice.

We also evaluated endosome adaptor levels in 9 month-old sciatic nerves to determine whether any clear patterns could be linked to the transport disruption. None of the significant changes identified in 3 month-old nerves were replicated in the older mice and fewer changes were observed in homozygous mutants at the later time point (**Figure 7**). At 9 months, *Yars*^*E196K/+*^ mice displayed reduced expression of dSnapin and Hook1, whereas *Yars*^*E196K/E196K*^ showed higher levels of dSnapin and lower levels of Hook1 (**Figure 7C-D**).

**Figure 7.**
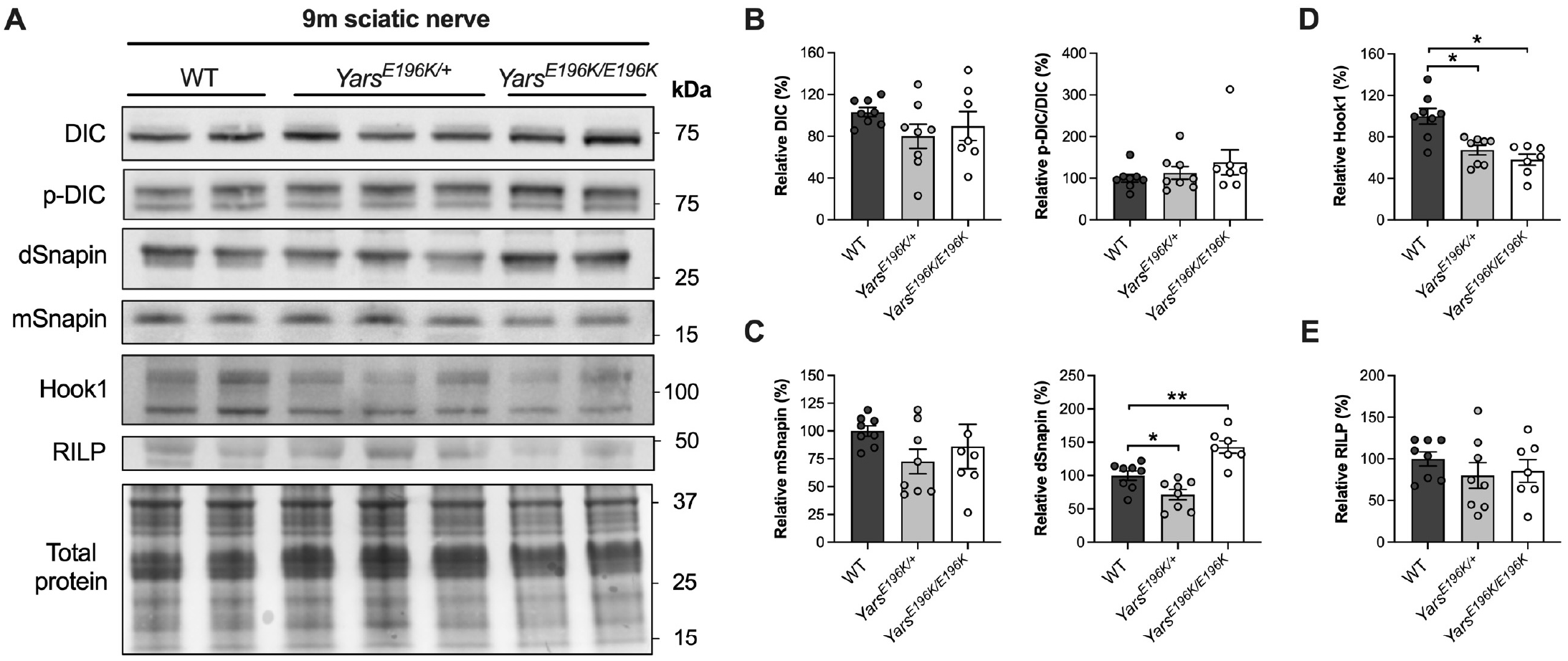
Endosome adaptor levels are altered in 9 month-old *Yars*^*E196K*^ sciatic nerves. (**A**) Western blots of dynein intermediate chain (DIC), phosphorylated DIC (p-DIC) and dynein adaptor proteins Snapin, Hook1 and RILP in sciatic nerves from 9 month-old wild-type, *Yars*^*E196K/+*^ and *Yars*^*E196K/E196K*^ mice. *kDa*, kilodalton. (**B-E**) Densitometric analyses of DIC (B, left, *P* = 0.298), p-DIC relative to DIC (B, right, *P* = 0.541 Kruskal-Wallis test), monomeric Snapin (mSnapin, C, left, *P* = 0.320), dimeric Snapin (dSnapin, C, right, *P* < 0.001), Hook1 (D, *P* = 0.003 Kruskal-Wallis test) and RILP (E, *P* = 0.517). The level of dSnapin is reduced in *Yars*^*E196K/+*^ and increased in *Yars*^*E196K/E196K*^, and the level of Hook1 is reduced in both *Yars*^*E196K*^ genotypes. For all graphs, genotypes were compared using one-way ANOVAs, unless otherwise stated; *n* = 7-8; \**P* < 0.05, \*\**P* < 0.01 Šídák’s/Dunn’s multiple comparisons test; means ± SEM plotted.

### Injection of TyrRS^E196K^ into wild-type muscle impairs endosome axonal transport

We have previously shown that injection of recombinant mutant GlyRS into wild-type muscles is sufficient to trigger non-cell autonomous impairment of endosome transport in otherwise healthy motor axons and that providing excess BDNF is able to overcome this disruption [76]. As mutant TyrRS aberrantly associates with the ECD of TrkB, we hypothesised that this mis-interaction at the nerve-muscle interface may be impairing BDNF-TrkB signalling and thus driving the trafficking defect observed in *Yars*^*E196K/E196K*^ mice. To test this idea, we bilaterally co-injected H_C_T with TyrRS into the tibialis anterior muscles of wild-type mice, with one side receiving TyrRS^WT^ and the other TyrRS^E196K^ (**Figure 8**). TyrRS^WT^ had no impact on signalling endosome axonal transport compared to vehicle-treated mice, whereas TyrRS^E196K^ caused a slow-down in endosome transport speeds compared to both vehicle and TyrRS^WT^ (**Figure 8A-B**), without observable differences in pausing (**Figure 8C-D**). These data indicate that DI-CMTC-causing mutant TyrRS within muscles is sufficient to non-cell autonomously impair signalling endosome axonal transport in wild-type mice.

**Figure 8.**
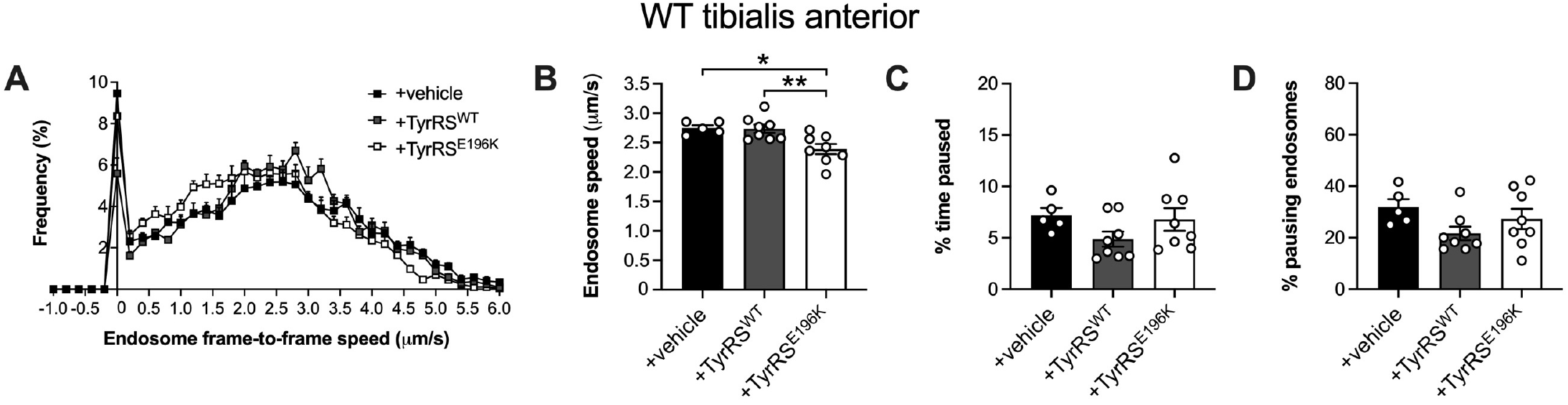
Exposure of wild-type motor terminals to TyrRS^E196K^, but not TyrRS^WT^, impairs *in vivo* axonal transport. (**A**) Endosome frame-to-frame speed histograms of 2-4-month-old wild-type (WT) mice receiving either unilateral tibialis anterior injections of PBS or bilateral injections of recombinant human TyrRS^WT^ into one tibialis anterior and TyrRS^E196K^ into the contralateral tibialis anterior. (**B-D**) TyrRS^E196K^ causes a slowdown in signalling endosome speed in otherwise healthy motor neurons (B, *P* = 0.004) without impacting the percentage of time paused (C, *P* = 0.193) or the percentage of pausing endosomes (D, *P* = 0.153). For all graphs, genotypes were compared using one-way ANOVAs; *n* = 5-8; \**P* < 0.05, \*\**P* < 0.01 Šídák’s multiple comparisons test; means ± SEM plotted.

To assess a possible relationship between the binding capacity of mutant ARS proteins and their effect on signalling endosome transport, we calculated the endosome transport speeds in wild-type mice receiving intramuscular injections of mutant GlyRS and mutant TyrRS relative to vehicle-treated controls (*n*.*b*., not raw speeds). Injection of GlyRS^L129P^ and GlyRS^G240R^ caused endosomes to be transported at 75.5 ± 3.6% and 75.7 ± 3.4% of control, respectively [76], whereas TyrRS^E196K^ resulted in transport at 87.0 ± 3.1% of vehicle-injected mice. Mutant forms of GlyRS thus cause a greater impact on transport than TyrRS^E196K^ (versus GlyRS^L129P^ *P* = 0.037 and versus GlyRS^G240R^ *P* = 0.038, unpaired *t*-tests), meaning that the axonal transport disruption caused by mutant ARS proteins correlates with the extent of their aberrant interaction with the ECD of TrkB.

### *Augmenting BDNF levels in muscles of* Yars^E196K/E196K^ *mice rescues axonal transport*

As *Yars*^*E196K/E196K*^ mice display a muscle-selective and age-dependent disruption in transport, similar in nature to that observed in CMT2D mice [76], we wished to establish whether increasing the availability of BDNF could also alleviate the trafficking disruption in this DI-CMTC model. To do so, we harnessed two different strategies – an acute paradigm with recombinant BDNF and a longer-term approach with AAV8-mediated, muscle-selective BDNF expression (**Figure 9A**).

**Figure 9.**
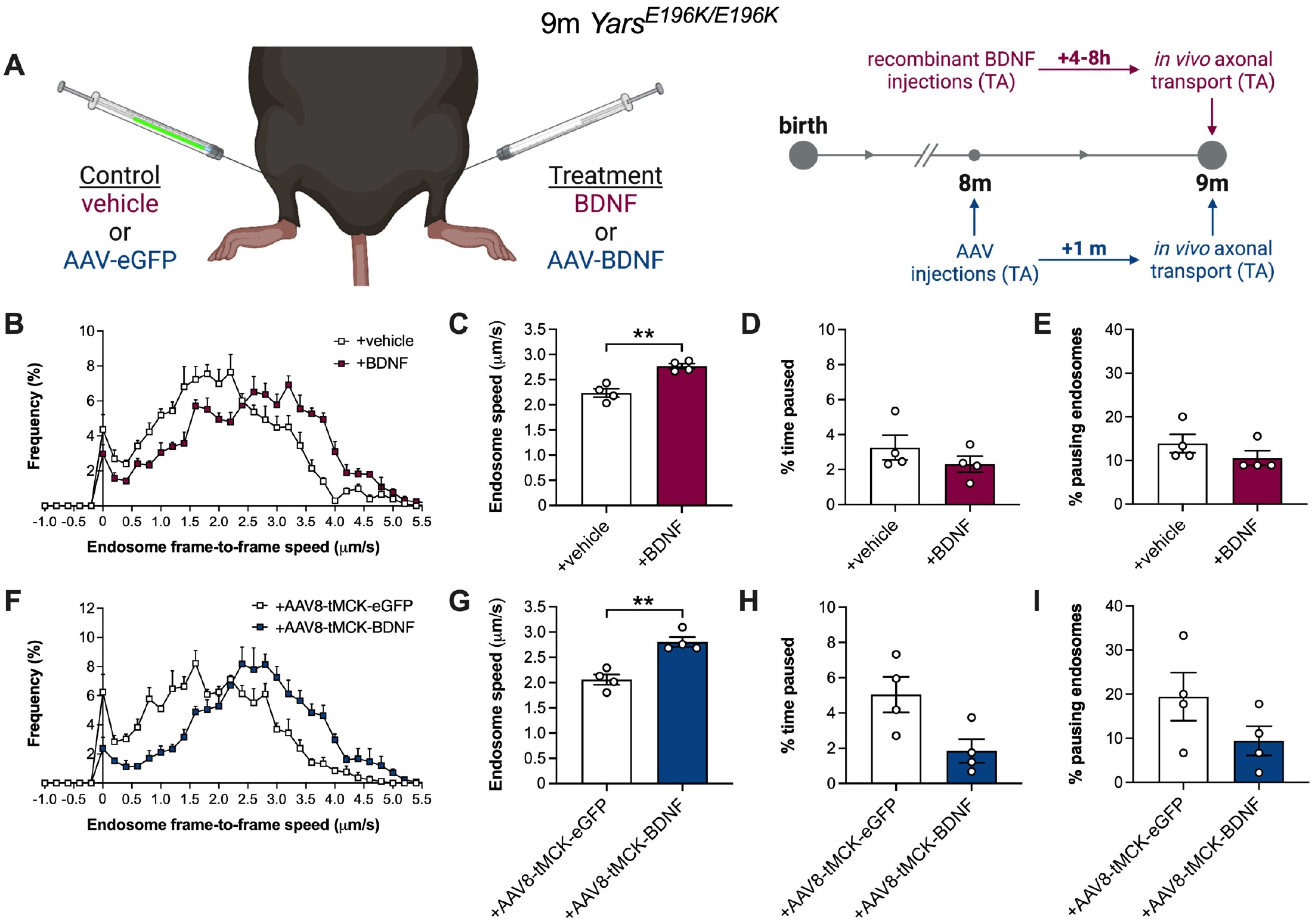
Boosting BDNF in muscles of homozygous *Yars*^*E196K*^ mice rescues *in vivo* axonal transport. (**A**) Schematics depicting the dual-tibialis anterior (TA) injection paradigms (left) and timelines (right) used in experiments to assess the impact of BDNF treatment in *Yars*^*E196K/E196K*^ mice. Animals either received dual injections of vehicle control versus recombinant BDNF at 9 months (maroon, top) or dual injections of AAV8-tMCK-eGFP versus AAV8-tMCK-BDNF at 8 months (blue, bottom). *In vivo* axonal transport of signalling endosomes was evaluated in tibialis anterior-innervating motor neurons at 9 months. Figure created using https://www.biorender.com. (**B**) Frame-to-frame speed histogram of signalling endosomes being transported within motor neurons innervating the tibialis anterior muscle of 9 month-old *Yars*^*E196K/E196K*^ mice 4-8 h post-treatment with intramuscular injections of vehicle or 25 ng recombinant BDNF. (**C-E**) BDNF increases the *in vivo* axonal transport speed of signalling endosomes in *Yars*^*E196K/E196K*^ (C, \*\**P* = 0.004), without impacting the percentage of time paused (D, *P* = 0.324) or the percentage of pausing endosomes (E, *P* = 0.625 Wilcoxon matched-pairs signed rank test). (**F**) Frame-to-frame speed histogram of signalling endosomes being transported within motor neurons innervating the tibialis anterior muscle of 9 month-old *Yars*^*E196K/E196K*^ mice 29-30 days after bilateral injection of AAV8-tMCK-eGFP into one tibialis anterior and AAV8-tMCK-BDNF into the contralateral tibialis anterior. (**G-I**) AAV8-tMCK-BDNF increases signalling endosome speed in *Yars*^*E196K/E196K*^ (G, \**P* = 0.032), without impacting the percentage of time paused (H, *P* = 0.116) or the percentage of pausing endosomes (I, *P* = 0.299). For all graphs, treatments were compared using paired *t*-tests, unless otherwise stated; *n* = 4; means ± SEM plotted. See also **Figure S10** and **Figure S11**.

We first evaluated the short-term effect of injecting recombinant human mature BDNF into muscles of 9 month-old *Yars*^*E196K/E196K*^ mice. This was done by co-injecting PBS vehicle control plus fluorescent H_C_T into one tibialis anterior muscle, and BDNF together with H_C_T into the contralateral muscle; signalling endosome transport was then sequentially assessed in both sciatic nerves 4-8 h-post injection. BDNF treatment caused an overt increase in the speed of signalling endosomes without significantly affecting pausing (**Figure 9B-E**), indicating that intramuscular injection of this neurotrophin is capable of rescuing the trafficking deficit. The endosome transport dynamics of vehicle-treated mice were similar to those previously identified in tibialis anterior-innervating axons of 9 month-old *Yars*^*E196K/E196K*^ mice (average speed: untreated 2.16 ± 0.1 µm/s versus vehicle-treated 2.24 ± 0.1 µm/s, *P* = 0.615 unpaired *t*-test).

The impact of longer-term exposure of *Yars*^*E196K/E196K*^ muscles to increased BDNF availability was analysed by injecting 8 month-old mice with AAV8-tMCK-eGFP control into one tibialis anterior and AAV8-tMCK-BDNF into the contralateral muscle. We have previously confirmed that the combination of muscle-tropic AAV serotype 8 [93] with the muscle-specific promoter tMCK [92] drives a robust increase in BDNF expression in muscles up to at least 30 days *in vivo* [76]. We therefore adapted this approach and treated 8 month-old *Yars*^*E196K/E196K*^ mice with bilateral injections of AAV into tibialis anterior muscles, before assessing transport of signalling endosomes at 9 months. We first confirmed that AAV treatment did indeed drive transgene expression in tibialis anterior muscles (**Figure S10**). We then showed that treatment with AAV8-tMCK-BDNF caused a strong increase in signalling endosome transport speed (**Figure 9F-G**), without significantly altering pausing (**Figure 9H-I**). Once again, we confirmed that the control virus had no impact on transport by comparing AAV8-tMCK-eGFP-treated mice with the original tibialis anterior-innervating axon data from 9 month-old *Yars*^*E196K/E196K*^ mice shown in **Figure 6** (average speed: untreated 2.16 ± 0.1 µm/s versus eGFP-treated 2.06 ± 0.1 µm/s, *P* = 0.562 unpaired *t*-test). This indicates that treatment with AAV8-tMCK-BDNF has a local effect within the injected muscle and does not systemically affect transport.

Given the female-specific reduction in *Yars*^*E196K/E196K*^ body weight (**Figure S5**), we compared transport in tibialis anterior-innervating neurons between 9 month-old *Yars*^*E196K/E196K*^ females and males, grouping all mice not receiving BDNF treatment (*i*.*e*., incorporating PBS vehicle- and AAV8-tMCK-eGFP-treated mice into the original 9 month data set). There was no difference in endosome trafficking between females and males (**Figure S11A-D**), suggesting that sex-specific differences are restricted to body weight and that transport in both sexes is equally defective.

Finally, to better evaluate whether treatment with BDNF is able to impact endosome pausing, we combined control-treated and BDNF-treated data from the acute and long-term transport experiments. Doing so revealed that boosting intramuscular levels of BDNF has a clear rescue effect on the axonal transport disruption in DI-CMTC mice (**Figure S11E-H**) – signalling endosome transport speeds were increased almost to wild-type levels, and both percentage of time paused and the percentage of pausing endosomes were reduced.

## Discussion

Homozygous *Yars*^*E196K*^ mice have previously been validated as a useful model for DI-CMTC, showing reduced NCVs, diminished motor axon calibres and a decline in motor endurance [36]. Here, we extend the temporal evaluation of *Yars*^*E196K*^ mice by assessing alternative features of peripheral nerve morphology and function at both 3 and 9 months. We identified several additional phenotypes pertinent to the human neuropathy, including distorted sensory neuron subpopulations, reduced motor neuron cell body areas, altered endosome adaptor levels within sciatic nerves, and impaired *in vivo* axonal transport of signalling endosomes (see **Figure S12** for a summary of the DI-CMTC phenotypes observed to date). This study provides the first evidence for axonal transport dysfunction in DI-CMTC and only the second study in which axonal transport has been identified as being impaired in mammalian peripheral neuropathy *in vivo*.

We also confirmed that motor neurons of *Yars*^*E196K/E196K*^ mice on the C57BL/6J background display a similar activation of the ISR as those mutants on C57BL/6N [79], which indicates that the subtle change in genetic background is unlikely to have impacted the overall severity of these mice. This is important, because we did not observe reduction in NMJ innervation across four different muscles nor any decline in maximal grip strength; thus, previously reported deficits in wire hang endurance are unlikely to result from overt anatomical changes at the neuromuscular synapse, but instead could reflect functional changes in synaptic physiology, similar to those identified in CMT2D mice [80]. The decline in motor neuron area is concordant with the previously identified smaller calibres of motor axons, which likely contribute to the slower NCVs and reduced motor endurance without impacting NMJ innervation and maximum muscle force. Indeed, reduced NCV occurs without denervation in several mouse models of neuropathy [4, 19, 45]. Nonetheless, we might have overlooked subtle morphological alterations at the neuromuscular synapse (e.g., reduced post-synaptic area or complexity), or by calculating the percentage of NMJs displaying degeneration, we might have missed denervated NMJs that have been rapidly lost from the muscle [18]. Alternatively, NMJ pathology may be overt in other muscles (e.g., the tibialis anterior).

The only clear phenotype that was present at 9 months, but not 3 months, was the axonal transport impairment, suggesting that the trafficking disruption in DI-CMTC is progressive, and therefore similar in nature to that identified in mutant *Gars* mice [76]. Disturbances in axonal transport have been identified in models of many different forms of both genetic and acquired peripheral neuropathy [5, 55, 73], including several different cellular and *in vivo* models of CMT2D [6, 43, 76, 78]; however, whether these deficits are a primary cause of disease or simply a secondary consequence of neuropathology remains an open question. In the case of DI-CMTC, given the temporal profile of peripheral nerve phenotypes in our study and the Hines *et al*. manuscript [36], it would appear that the endosome trafficking deficits are a secondary feature of disease. Nevertheless, they represent a primary effect of mutant TyrRS and are likely to exacerbate pathology by reducing the long-range, pro-survival neurotrophin signalling from motor nerve terminals to the spinal cord. It remains to be determined at what stage the transport becomes impaired and whether assessments beyond 9 months will reveal additional phenotypes subsequent to the transport disruption, such as loss of NMJ integrity.

The relevance of changes in endosome motor adaptor protein levels in 3 and 9 month sciatic nerves of *Yars*^*E196K*^ mice remains unclear, as we did not see consistent patterns correlating with age and/or the development of axonal transport disruption. A possible explanation for this is that different motor adaptor proteins may be required for signalling endosome trafficking as they progress from distal to proximal segments of the axon [15]. Indeed, the retrograde delivery of signalling endosomes in motor neurons relies on Rab7 [21], whereas different effectors mediate the retrograde transport and maturation of autophagic organelles [14]. Therefore, impairments in BDNF-TrkB signalling at the axon terminal caused by mutant TyrRS interacting with TrkB may have multifarious age- and location-dependent effects on the endosome motor adaptors, obscuring a clear neuropathic signature. That being said, the three endosome adaptors that were increased at 3 months in *Yars*^*E196K/E196K*^ nerves were no longer elevated at 9 months, perhaps indicative of a possible loss of compensatory mechanisms with age.

Replicating findings obtained with mutant GlyRS, we show that mutant TyrRS aberrantly associates with the ECD of the BDNF receptor TrkB and that injection of TyrRS^E196K^ into wild-type muscle is sufficient to impair *in vivo* axonal transport of signalling endosomes in otherwise healthy motor axons. We have previously determined that inhibition of the BDNF-TrkB signalling pathway in wild-type muscles, either pharmacologically or using BDNF blocking antibodies, results in impaired signalling endosome trafficking [76]. It is thus conceivable that TyrRS^E196K^ is mis-interacting with TrkB at motor nerve terminals, impinging upon BDNF signalling and slowing down endosome trafficking. Consistent with this hypothesis, provision of excess BDNF in DI-CMTC muscles fully corrects the transport phenotype. As we did not detect deficits in this dynamic process in heterozygous *Yars*^*E196K*^ mice even at 15 months, it is likely to be a dose-dependent disruption. Consistent with this idea, mutant GlyRS, which shows a stronger association with the ECD of TrkB, causes a greater transport deficit when injected into wild-type mouse muscles, and CMT2D mice display an earlier and more severe trafficking defect.

The interaction between TyrRS^E196K^ and TrkB is likely to impair the ability of BDNF to signal through TrkB, which could result in the slowing of neurotrophin-containing signalling endosomes via several different, non-mutually exclusive mechanisms, including reduced transcription, translation, phosphorylation and/or recruitment of adaptor and motor proteins [42, 61, 89, 90]. We are currently working to understand the molecular mechanism underpinning the slowed transport and have previously discussed these possibilities [76], so will not elaborate further here. It remains unknown why the TyrRS^E196K^ dosage does not differentially affect the sensory neuron populations in lumbar DRG of *Yars*^*E196K*^ mice; however, the data suggest that, since this is a toxic gain-of-function, once a maximum threshold level of mutant TyrRS is available (such as that reached in the heterozygous mutants), no further changes to sensory neuron populations occur.

Mutations in *YARS1* enhance the binding partners TRIM28 and HDAC1 [9], as well as F-actin [23]. In this study, TyrRS^WT^ also appears to interact with TrkB at a low level compared with mutant TyrRS, whereas previous data indicate that this is not the case for GlyRS^WT^ [68, 76]. Therefore, it is possible that TyrRS^WT^ constitutively interacts with TrkB to exert a non-canonical function, rather than being a novel aberrant association, but further work is required to address this.

As aminoacyl-tRNA synthetases represent the largest protein family linked to CMT aetiology, it is an alluring prospect that they share a common pathomechanism, which can be targeted to treat the several different associated subtypes of CMTs. Supporting this possibility, mouse and fly models of CMT2D and DI-CMTC display protein synthesis impairments in motor neurons linked to the ISR, which are driven by tRNA^Gly^ sequestration and ribosome stalling in mutant *Gars* models [47, 79, 97]. Moreover, co-genetic modifiers of pathology have been discovered in *Drosophila* models of *GARS1*- and *YARS1*-associated neuropathy [24]. Our work contributes another shared mechanism between CMT2D and DI-CMTC that causes impairment in signalling endosome axonal transport through perturbations of the BDNF-TrkB signalling axis. There is scope for this pathomechanism to be relevant to additional ARS-linked neuropathies, since structural relaxation and conformational opening have been reported for both CMT-linked alanyl-tRNA synthetase [82] and histidyl-tRNA synthetase [11] mutants. Mammalian models for other subtypes of ARS-linked neuropathy are not currently available to determine whether axonal transport disruption is impaired in a mammalian setting *in vivo*, although with the advent of CRISPR/Cas9 technology, they are likely forthcoming. While the interruption in cargo trafficking within axons may not be the primary cause of neuropathy linked to ARS mutations, it will certainly contribute to the demise of peripheral nerves and is a promising candidate process for pharmacological targeting [26, 32].

Through acute and month-long treatment paradigms, we have shown that boosting the availability of BDNF at motor nerve terminals of *Yars*^*E196K/E196K*^ mice rescues deficits in axonal transport of neurotrophin-containing signalling endosomes *in vivo*, replicating findings in CMT2D mice [76]. Targeting muscle may therefore provide an effective approach to treating ARS-linked neuropathies, although care must be taken with concentration, timing and duration of neurotrophic factor delivery to ensure the best chance of treatment success [22]. In combination with additional interventions that have proven beneficial in mutant *Gars* mice, including HDAC6 inhibition [6, 43, 78], delivery of neurotrophin 3 [51] and VEGF_165_ [34], and provision of excess tRNA^Gly^ [97], boosting BDNF in muscles may serve as part of a combinatorial treatment to address the therapeutic needs of one of the most common inherited neuromuscular conditions.

## Author Contributions

Conceptualisation: JNS. Investigation: ERR, RLS, JQ, DV-C, SS, YT, RS, JNS. Writing and figure production: JNS with input from all authors. Contributed *Yars*^*E196K*^ mice: RWB. Supervision and funding: RWB, XLY, GS, JNS.

## Materials & Correspondence

AAV expression plasmids are covered by an MTA with OXGENE (UK). Material requests should be addressed to the corresponding authors.

## Acknowledgements

We thank Timothy J. Hines (Jackson Laboratory) for sharing the sequences of *Yars*^*E196K*^ genotyping primers, K. Kevin Pfister (University of Virginia) for donating the p-DIC (S81) antibody, and personnel of the Denny Brown Laboratory (UCL) for assistance with mouse colonies.

## Funding

This project was funded by Medical Research Council awards MR/S006990/1 and MR/Y010949/1 (JNS); a Rosetrees Trust grant M806 (JNS, GS); the UCL Neurogenetic Therapies Programme funded by The Sigrid Rausing Trust (JNS, GS); the UCL Therapeutic Acceleration Support scheme supported by funding from MRC IAA 2021 UCL MR/X502984/1 (JNS); a Human Frontier Science Program Long-Term Fellowship LT000220/2017-L (SS); National Institutes of Health awards R37 NS05415 (RWB) and R35 GM139627 (XLY); Wellcome Trust awards 107116/Z/15/Z and 223022/Z/21/Z (GS); and a UK Dementia Research Institute award (UK DRI-1005) through UK DRI Ltd, principally funded by the UK Medical Research Council (GS).

## Conflict of Interest Statement

The technology described in this work has been protected in the patent GB2303495.2 (patent applicant, UCL Business Ltd., status pending), in which GS and JNS are named as inventors. The other authors declare no competing interests.

## Supplementary Information

### Supplementary Figures

**Supplementary Figure S1.**
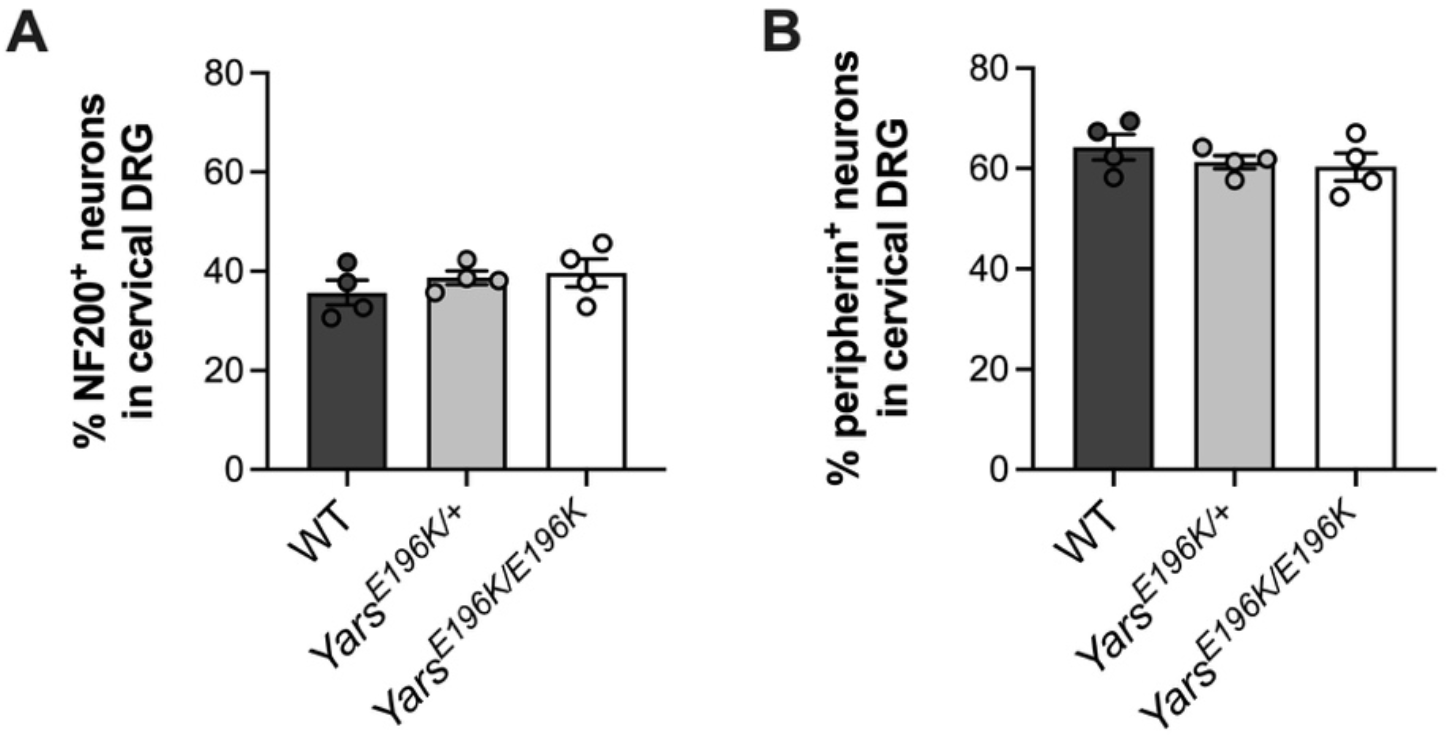
Sensory neuron subtypes are unaffected in *Yars*^*E196K*^ cervical DRG. (**A-B**) There is no difference in the percentage of NF200^+^ (A, *P* = 0.475) or peripherin^+^ (B, *P* = 0.475) neurons in cervical DRG between wild-type and *Yars*^*E196K*^ mice. In both graphs, genotypes were compared using one-way ANOVAs; *n* = 4; means ± standard error of the mean (SEM) plotted.

**Supplementary Figure S2.**
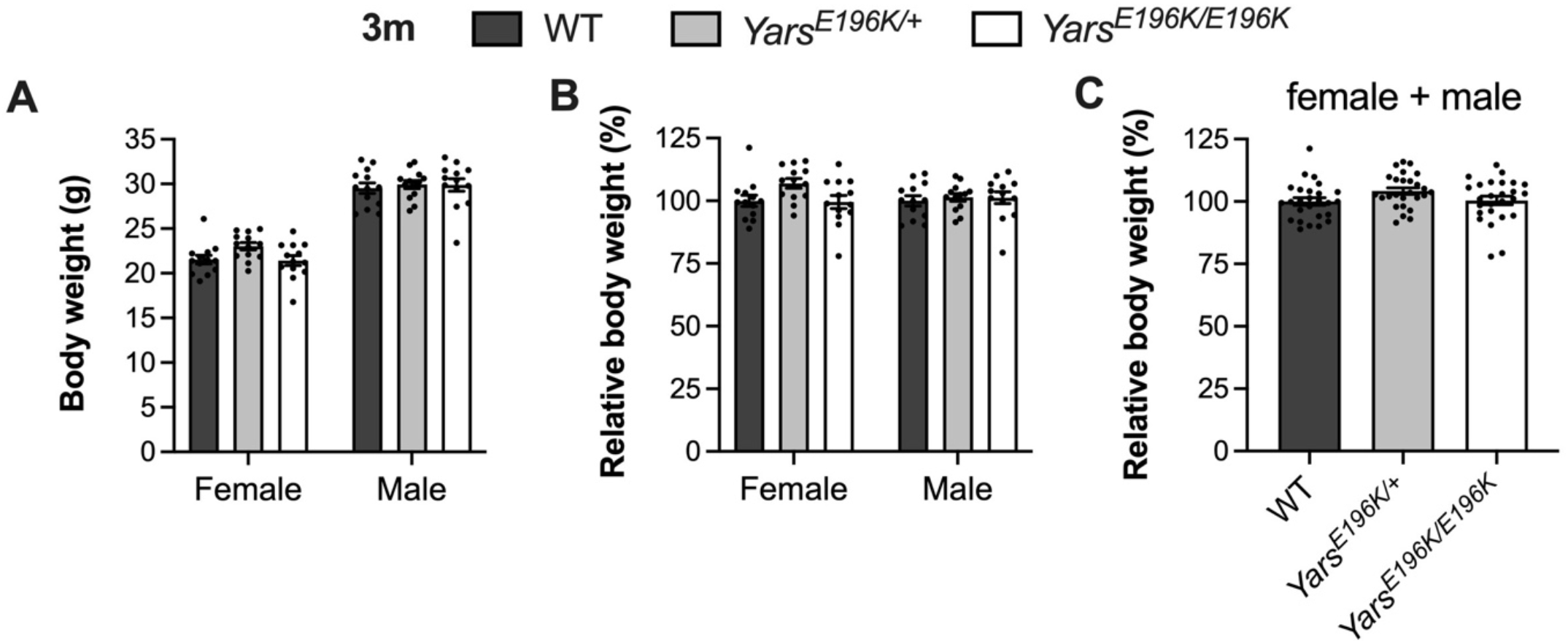
*Yars*^*E196K*^ females and males display normal body weight at 3 months. (**A-B**) There is no difference in body weight (A, genotype *P* = 0.169, sex *P* < 0.001, interaction *P* = 0.337 two-way ANOVA) or relative body weight (B, genotype *P* = 0.105, sex *P* = 0.457, interaction *P* = 0.208 two-way ANOVA) between female or male wild-type, *Yars*^*E196K/+*^ and *Yars*^*E196K/E196K*^ mice at 3 months. *n* = 13. (**C**) When relative grip strenths of females and males are combined, there is still no difference between genotypes (*P* = 0.107 one-way ANOVA). *n* = 26. For all graphs, means ± SEM plotted.

**Supplementary Figure S3.**
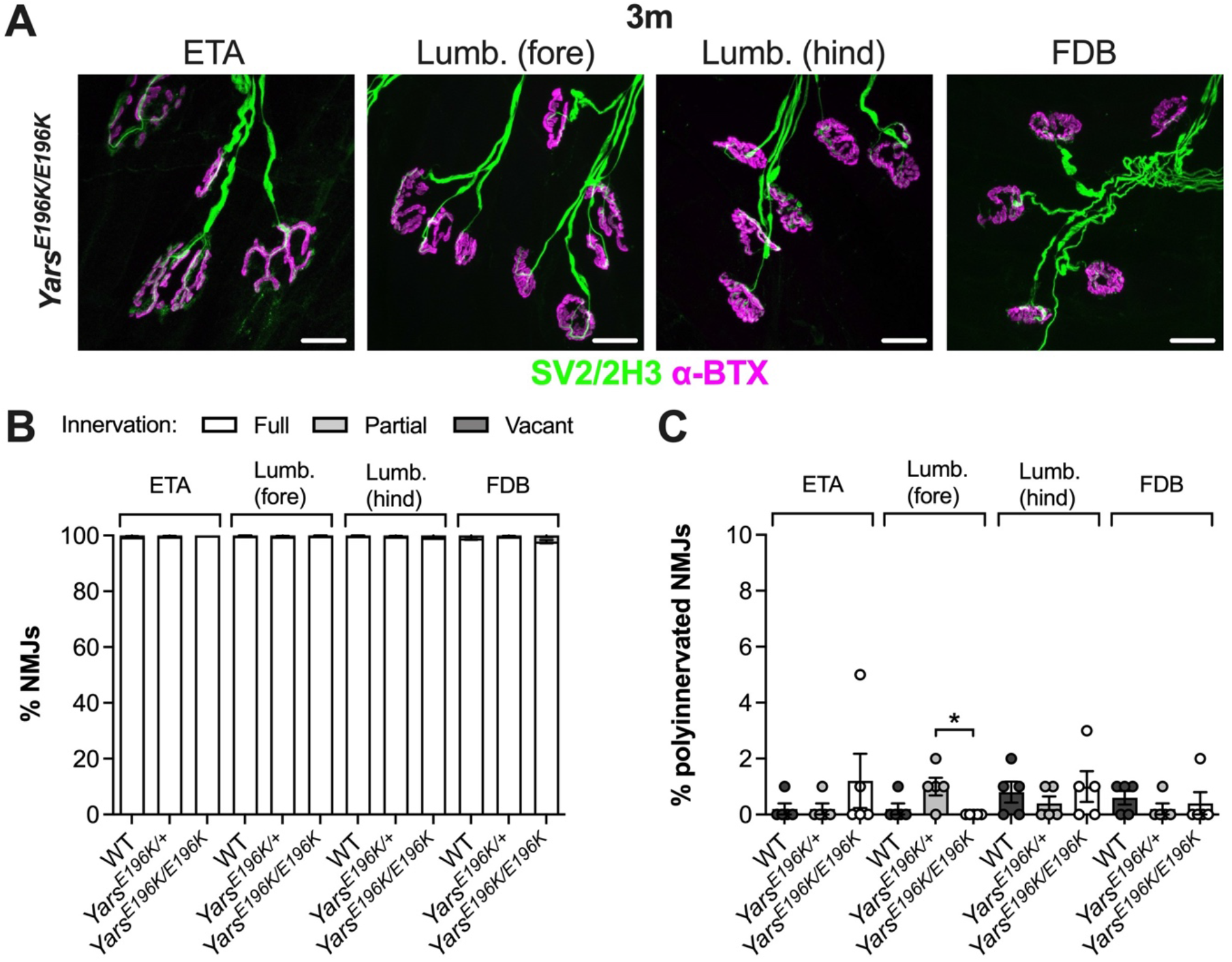
Neuromuscular junction innervation and maturation are unaffected in *Yars*^*E196K*^ mice at 3 months. (**A**) Representative collapsed z-stack confocal images of NMJs in ETA, forepaw lumbrical [*Lumb. (fore)*], hindpaw lumbrical [*Lumb. (hind)*] and FDB muscles in a 3 month-old *Yars*^*E196K/E196K*^ mouse. Lower motor neurons are visualised using a combination of SV2/2H3 (green) and post-synaptic AChRs with α-BTX (magenta). These images are also representative of wild-type and *Yars*^*E196K/+*^ mice. Scale bars = 20 µm. (**B**) No difference between genotypes in NMJ innervation was observed in the ETA (*P* = 0.251), the forepaw lumbricals (*P* > 0.999), hindpaw lumbricals (*P* = 0.501) or FDB (*P* = 0.063). (**C**) No meaningful difference between genotypes in the percentage of polyinnervated NMJs was observed in the ETA (*P* = 0.725), the forepaw lumbricals (*P* = 0.041), hindpaw lumbricals (*P* = 0.750) or FDB (*P* = 0.501). For all graphs, genotypes were compared using Kruskal-Wallis tests; *n* = 5; \**P* < 0.05 Dunn’s multiple comparisons test; means ± SEM plotted.

**Supplementary Figure S4.**
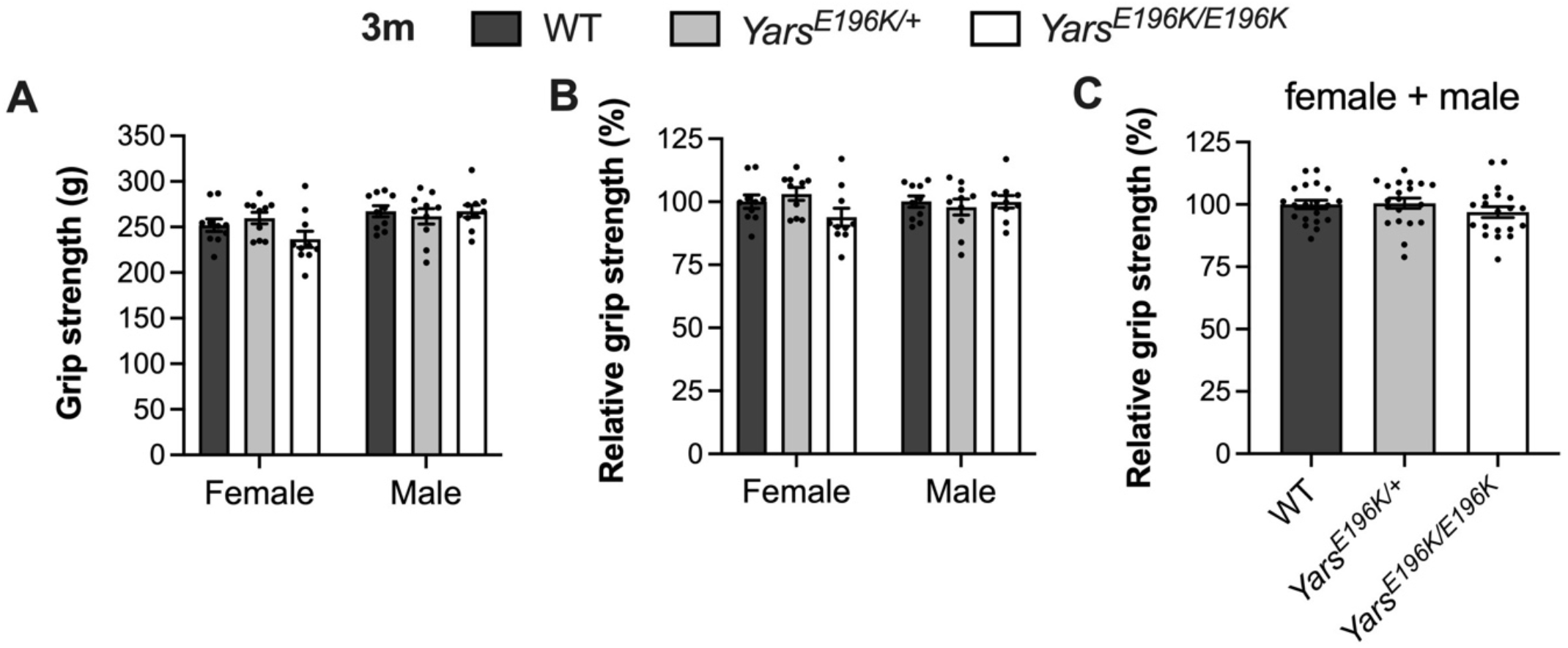
Grip strength of *Yars*^*E196K*^ mice is unaffected at 3 months. (**A-B**) There is no difference in grip strength (A, genotype *P* = 0.420, sex *P* < 0.010, interaction *P*= 0.150 two-way ANOVA) or relative grip strength (B, genotype *P* = 0.391, sex *P* = 0.896, interaction *P* = 0.141 two-way ANOVA) between female or male wild-type, *Yars*^*E196K/+*^ and *Yars*^*E196K/E196K*^ mice at 3 months. *n* = 10. (**C**) When relative body weights of females and males are combined, there is still no difference between genotypes (*P* = 0.398 one-way ANOVA). *N* = 20. For all graphs, means ± SEM plotted.

**Supplementary Figure S5.**
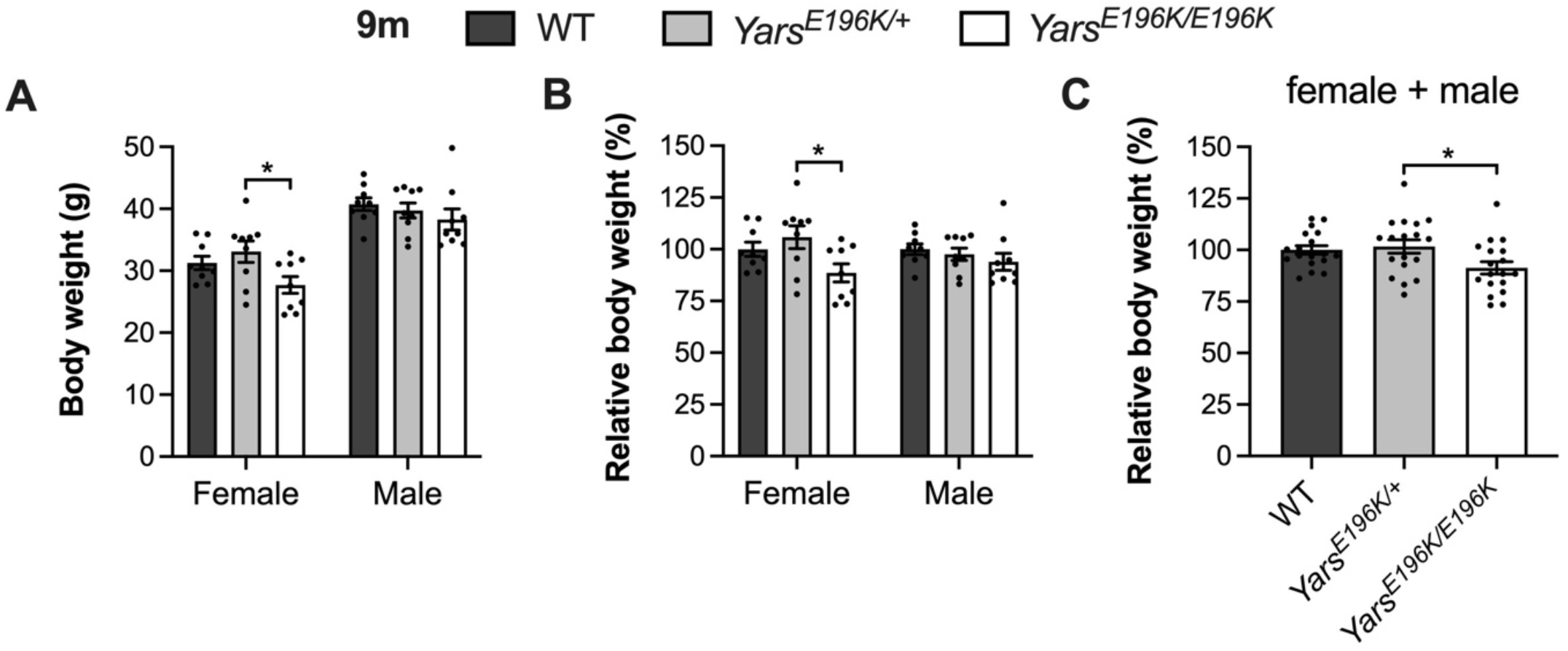
Homozygous *Yars*^*E196K*^ mice display a small reduction in body weight at 9 months. (**A-B**) 9 month-old female *Yars*^*E196K/E196K*^ mice display a reduction in body weight (A, genotype *P* = 0.033, sex *P* < 0.001, interaction *P* = 0.347 two-way ANOVA) and relative body weight (B, genotype *P* = 0.025, sex *P* = 0.764, interaction *P* = 0.230 two-way ANOVA) compared with *Yars*^*E196K/+*^. *n* = 9. (**C**) When relative body weights of females and males are combined, *Yars*^*E196K/E196K*^ mice display a deficit compared to *Yars*^*E196K/+*^, but not wild-type (*P* = 0.025 one-way ANOVA). *n* = 18. For all graphs, \**P* < 0.05 Šidák’s multiple comparisons test; means ± SEM plotted.

**Supplementary Figure S6.**
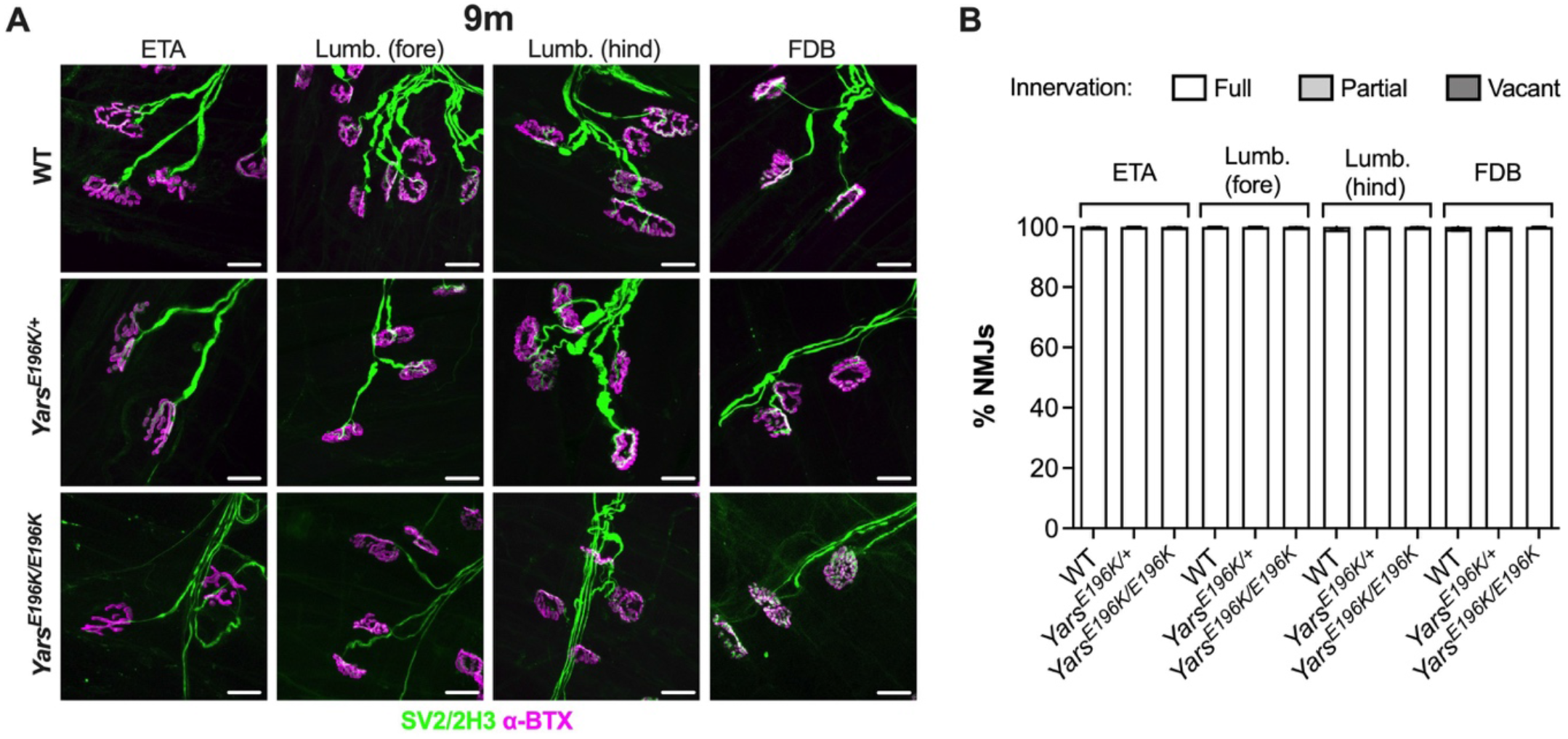
Neuromuscular junction innervation remains unaffected in *Yars*^*E196K*^ mice at 9 months. (**A**) Representative collapsed z-stack confocal images of NMJs in ETA, forepaw lumbrical [*Lumb. (fore)*], hindpaw lumbrical [*Lumb. (hind)*] and FDB muscles in 9 month-old wild-type, *Yars*^*E196K/+*^ and *Yars*^*E196K/E196K*^ mice. Lower motor neurons are visualised using a combination of SV2/2H3 (green) and post-synaptic AChRs with α-BTX (magenta). Scale bars = 20 µm. (**B**) No difference between genotypes in NMJ innervation was observed in the ETA (*P* > 0.999), the forepaw lumbricals (*P* > 0.999), hindpaw lumbricals (*P* = 0.442) or FDB (*P* = 0.297). Genotypes were compared using Kruskal-Wallis tests; *n* = 6; means ± SEM plotted.

**Supplementary Figure S7.**
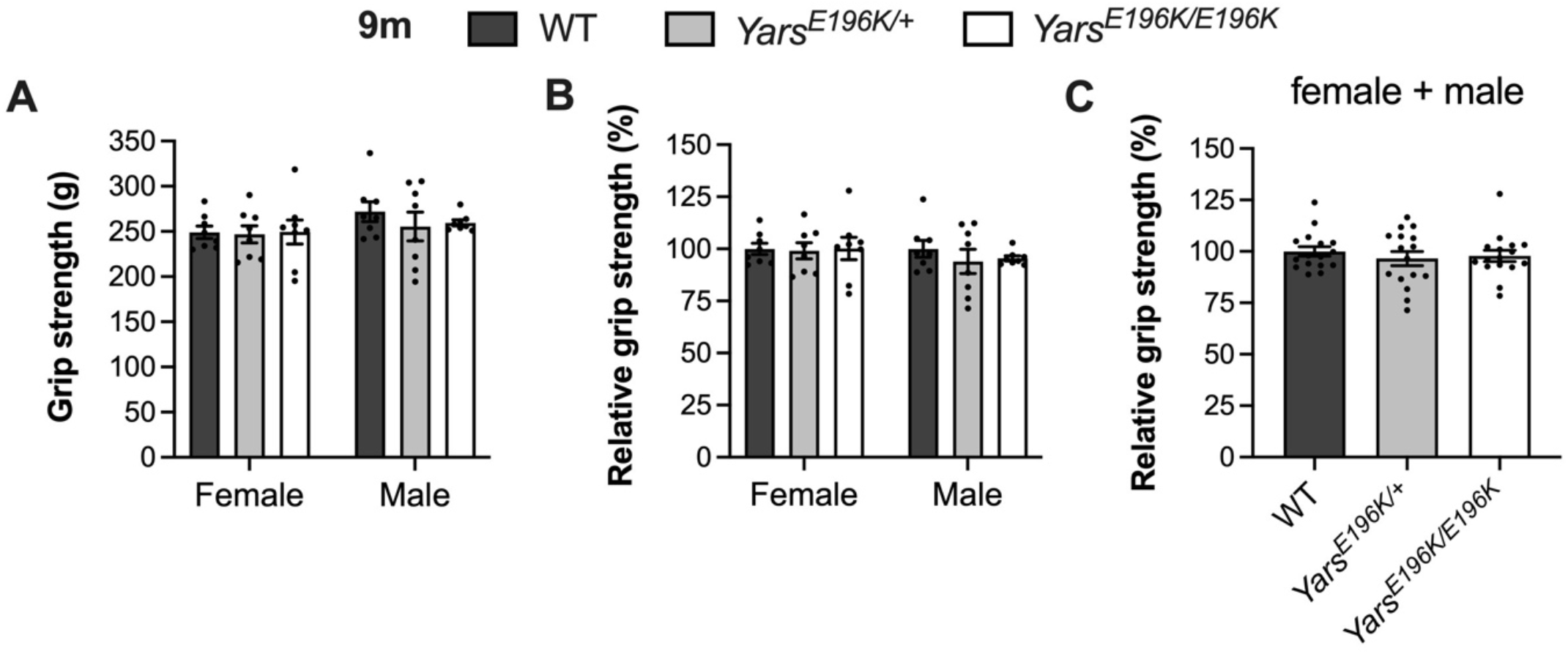
Grip strength of *Yars*^*E196K*^ mice is unaffected at 9 months. (**A-B**) There is no difference in grip strength (A, genotype *P* = 0.682, sex *P* = 0.122, interaction *P* = 0.773 two-way ANOVA) or relative grip strength (B, genotype *P* = 0.698, sex *P* = 0.340, interaction *P* = 0.792 two-way ANOVA) between female or male wild-type, *Yars*^*E196K/+*^ and *Yars*^*E196K/E196K*^ mice at 9 months. *n* = 8. (**C**) When relative grip strengths of females and males are combined, there is still no difference between genotypes (*P* = 0.689 one-way ANOVA). *N* = 16. For all graphs, means ± SEM plotted.

**Supplementary Figure S8.**
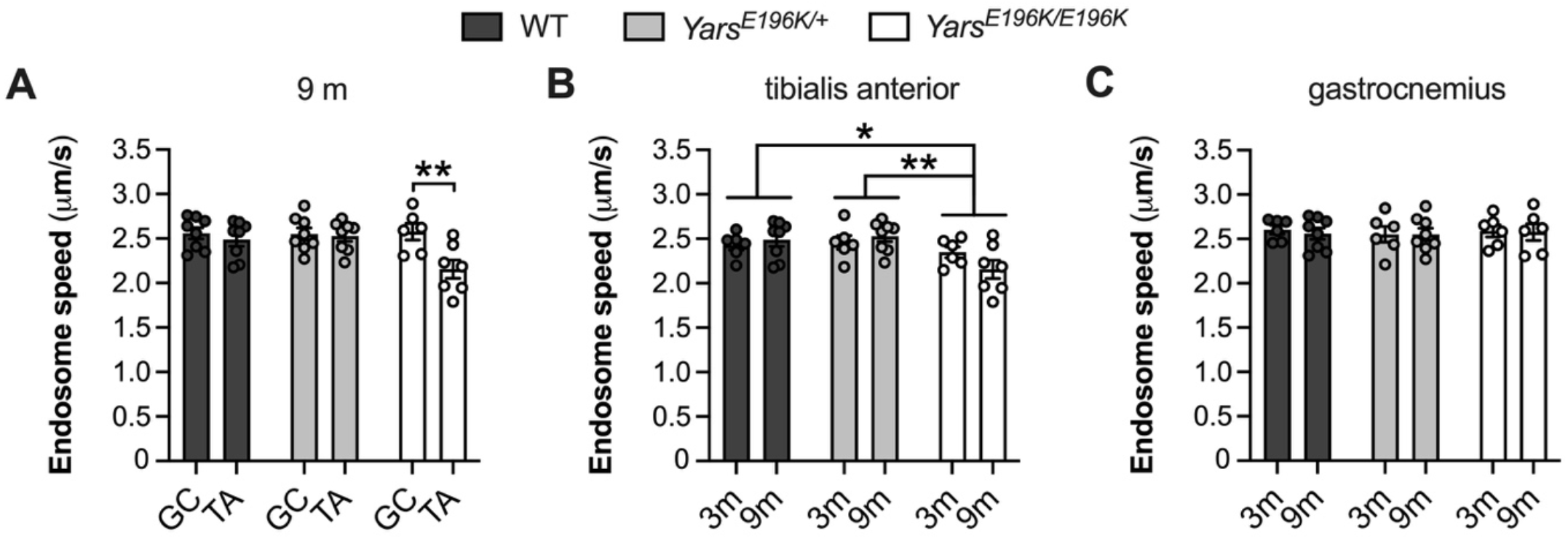
9 month-old *Yars*^*E196K*^ homozygotes display selective impairment in *in vivo* axonal transport of signaling endosomes. (**A**) There is a reduction in signaling endosome axonal transport speed in motor axons innervating the tibialis anterior (TA), but not gastrocnemius (GC), in *Yars*^*E196K/E196K*^ mice aged 9 months. No differences are observed between muscles in wild-type and *Yars*^*E196K/+*^ mice (genotype *P* = 0.068, muscle *P* < 0.010, interaction *P* = 0.033 two-way ANOVA). (**B-C**) Taking into account endosome transport speeds at 3 (3m) and 9 months (9m) of age, *Yars*^*E196K/E196K*^ mice display a reduction in the tibialis anterior (B, genotype *P* = 0.007, age *P* = 0.747, interaction *P* = 0.164 two-way ANOVA), but not gastrocnemius (C, genotype *P* = 0.888, age *P* = 0.752, interaction *P* = 0.953 two-way ANOVA), compared to wild-type and *Yars*^*E196K/+*^. For all graphs, *n* = 6-8; \**P* < 0.05, \*\**P* < 0.01 Šidák’s multiple comparisons test; means ± SEM plotted. These data are also presented in **Figure 3** and **Figure 6**.

**Supplementary Figure S9.**
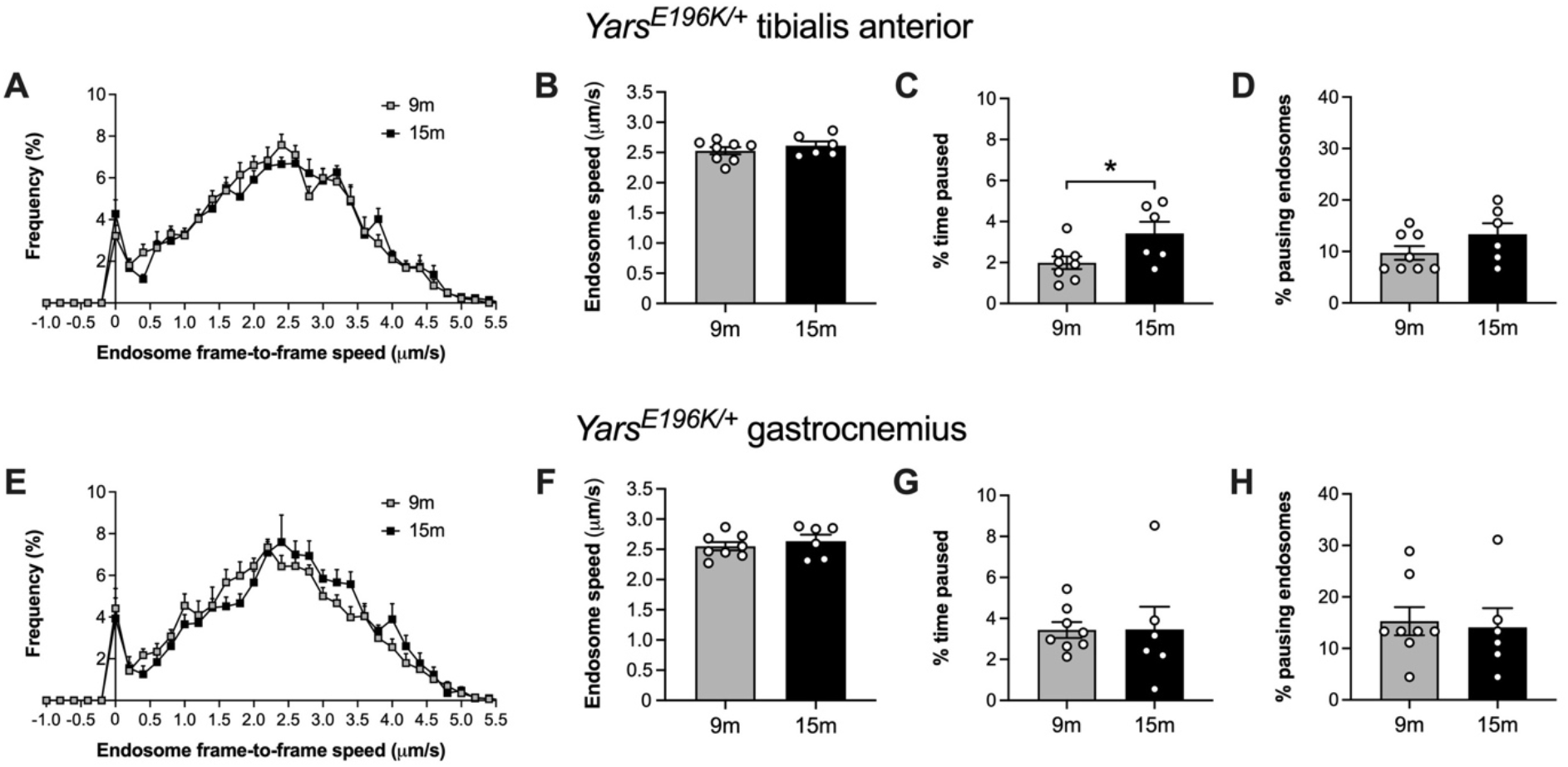
*In vivo* axonal transport remains largely unperturbed in *Yars*^*E196K*^ heterozygous mice at 15 months. (**A**) Frame-to-frame speed histogram of signalling endosomes being transported within motor neurons innervating the tibialis anterior muscle of *Yars*^*E196K/+*^ mice aged 9 (9m) and 15 months (15m). (**B-D**) In *Yars*^*E196K/+*^ neurons innervating the tibialis anterior, there is no difference between time points in signalling endosome speed (B, *P* = 0.380 unpaired *t*-test) or the percentage of pausing endosomes (D, *P* = 0.161 Mann-Whitney *U* test), but there is a small increase in percentage of time paused (C, * *P* = 0.036 unpaired *t*-test) at 15 months. (**E**) Frame-to-frame speed histogram of signalling endosomes being transported within motor neurons innervating the gastrocnemius muscle of *Yars*^*E196K/+*^ mice aged 9 and 15 months. (**F-H**) In *Yars*^*E196K/+*^ neurons innervating the gastrocnemius, there is no difference between time points in signalling endosome speed (F, *P* = 0.501 unpaired *t*-test), percentage of time paused (G, *P* = 0.979 unpaired *t*-test) or the percentage of pausing endosomes (H, *P* = 0.729 Mann-Whitney *U* test). For all graphs, *n* = 6-8; means ± SEM plotted. The 9m data are also presented in **Figure 6** and **Figure S8**.

**Supplementary Figure S10.**
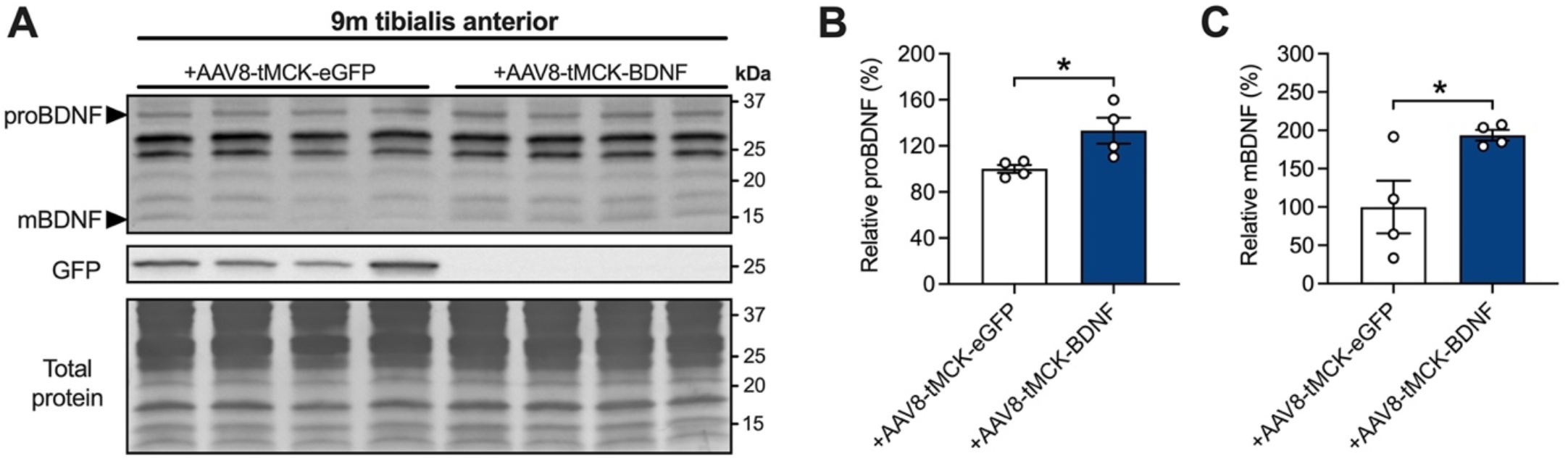
AAV8-tMCK drives transgene expression after intramuscular injection. (**A**) Western blots of GFP and BDNF in tibialis anterior muscles from 9 month-old *Yars*^*E196K/E196K*^ mice that received dual injections of AAV8-tMCK-eGFP versus AAV8-tMCK-BDNF at 8 months. GFP is expressed in muscles injected with AAV8-tMCK-eGFP. *kDa*, kilodalton. (**B-C**) Densitometric analyses of proBDNF (B) and mature BDNF (mBDNF, C) reveal increases in BDNF availability in muscles injected with AAV8-tMCK-BDNF. For both graphs, *n* = 4; \**P* < 0.05 unpaired *t*-test; means ± SEM plotted.

**Supplementary Figure S11.**
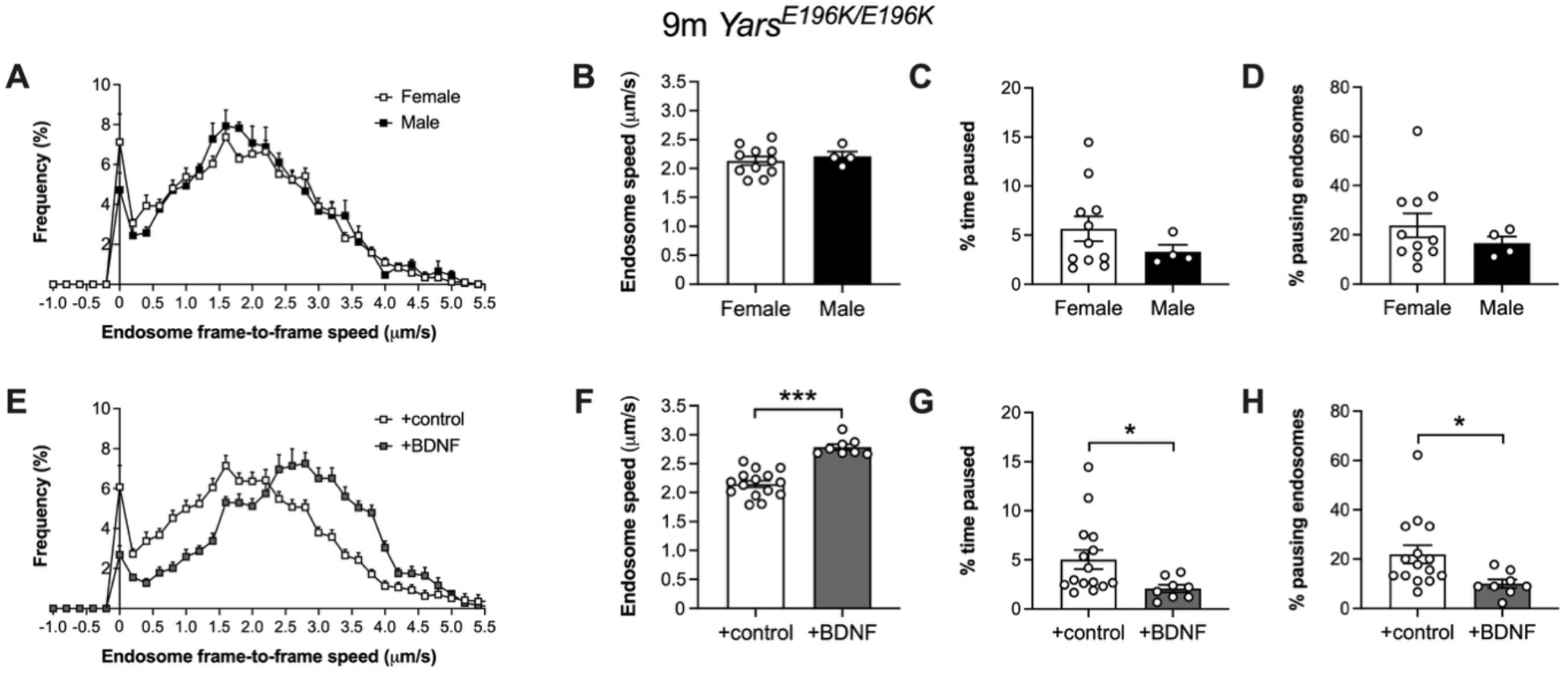
Boosting BDNF in muscles of *Yars*^*E196K*^ homozygotes rescues *in vivo* axonal transport of signaling endosomes. (**A**) Frame-to-frame speed histogram of signalling endosomes being transported within motor neurons innervating the tibialis anterior muscle of 9 month-old *Yars*^*E196K/E196K*^ females and males. (**B-D**) There is no difference between 9 month-old female and male *Yars*^*E196K/E196K*^ mice in signaling endosome transport dynamics in tibialis anterior-innervating motor axons. Endosome transport speeds (B, *P* = 0.579), percentage time paused (C, *P* = 0.306) and percentage pausing endosomes (D, *P* = 0.407) all show no difference between sexes. *n* = 4-11 (**E**) Frame-to-frame speed histogram of signalling endosomes being transported within motor neurons innervating the tibialis anterior muscle of 9 month-old *Yars*^*E196K/E196K*^ that were either untreated/control-treated (white) or received acute/AAV-mediated treatment with BDNF (grey). (**F-H**) Boosting BDNF in muscle increases signalling endosome transport speed (F, \*\*\**P* < 0.001), reduces the percentage of time paused (G, \**P* = 0.013) and diminishes the percentage of pausing endosomes (H, \**P* = 0.033). For all graphs, sexes/treatments were compared using unpaired *t*-tests; *n* = 4-11 (A-D) and 8-15 (E-H); means ± SEM plotted. These data are also presented in **Figure 6** and **Figure 9**.

**Supplementary Figure S12.**
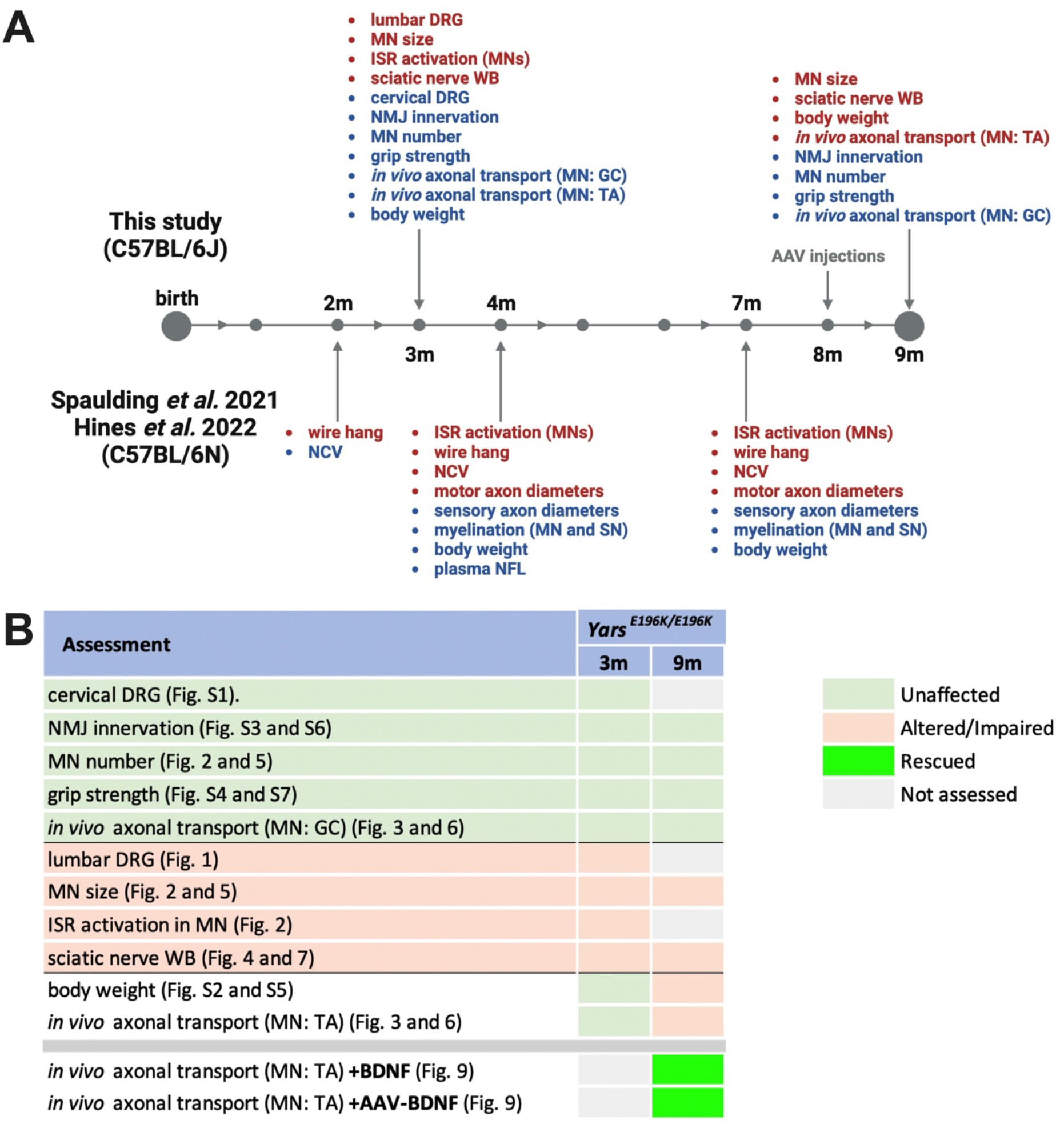
Summary of homozygous *Yars*^*E196K*^ mouse phenotypes. (**A**) Timeline of *Yars*^*E196K/E196K*^ phenotypes identified in this study (above timeline) and that of Spaulding *et al*. 2021 and Hines *et al*. 2022 (below timeline) [2, 4]. Assessments written in blue indicate that no phenotype was observed and those written in red identify a defect (compared to wild-type and/or *Yars*^*E196K/+*^). Figure created using https://www.biorender.com. (**B**) Summary table of homozygous *Yars*^*E196K*^ phenotypes identified in this study. At 3 months, *Yars*^*E196K/E196K*^ mice display 1) perturbed populations of sensory neuron subtypes in lumbar, but not cervical DRG, 2) reduced cell body size and increased ISR activation in lumbar motor neurons, and 3) altered levels of endosome adaptors in the sciatic nerve. No deficits in weight, NMJ innervation, grip strength or *in vivo* axonal transport of signaling endosomes were observed. 9 month-old *Yars*^*E196K/E196K*^ mice additionally display a mild, sex-dependent reduction in weight and selective impairment in *in vivo* axonal transport of signaling endosomes in motor axons innervating the tibialis anterior, but not gastrocnemius. Boosting BDNF in muscle of *Yars*^*E196K/E196K*^ mice, either through injection of recombinant BDNF or BDNF-producing AAV, is able to alleviate this transport dysfunction. For both panels, *GC*, gastrocnemius; *ISR*, integrated stress response; *MN*, motor neuron; *NCV*, nerve conduction velocity; *NFL*, Neurofilament light chain; *SN*, sensory neuron; *TA*, tibialis anterior; *WB*, western blot.

### Supplementary Tables

**Table S1.** Experimental sample sizes and mouse details. *F*, female; *GC*, gastrocnemius; *Het, Yars*^*E196K/+*^; *Hom, Yars*^*E196K/E196K*^; *M*, male; *m*, months; *MN*, motor neuron; *n*, sample size; *n*.*a*., not applicable; *TA*, tibialis anterior; *TyrRS*, tyrosyl-tRNA synthetase; *WT*, wild-type.

| Figure | Data | *n* | Sex (WT/Het/Hom) | Age (d) |
| --- | --- | --- | --- | --- |
| 1B | *in vitro* pull-downs | 3 | n.a. | n.a. |
| 1D | 3 m lumbar DRG neuron subtypes | 4 | 2M2F / 2M2F / 3M1F | 91-99 |
| S1 | 3 m cervical DRG neuron subtypes | 4 | 2M2F / 2M2F / 3M1F | 91-99 |
| S2 | 3 m body weight | 13/26 | 13M13F / 13M13F / 13M13F | 89-99 |
| S3 | 3 m NMJ phenotypes | 5 | 5M0F / 2M3F / 2M3F | 90-99 |
| 2 | 3 m spinal cord MNs | 4 | 2M2F / 3M1F / 2M2F | 91-99 |
| 3 | 3 m axonal transport in TA and GC | 6 | 5M1F / 6M0F / 3M3F | 91-97 |
| 4 | 3 m sciatic nerve blots | 4 | 4M0F / 2M2F / 2M2F | 90-99 |
| S4 | 3 m grip strength | 10/20 | 10M10F / 10M10F / 10M10F | 86-99 |
| S5 | 9 m body weight | 9/18 | 9M9F / 9M9F / 9M9F | 267-285 |
| S6 | 9 m NMJ phenotypes | 6 | 1M5F / 3M3F / 1M5F | 267-280 |
| 5 | 9 m spinal cord MN counts and areas | 4-7 | 4M0F / 4M1F / 5M0-2F | 271-281 |
| 6 | 9 m axonal transport in TA and GC | 6-8 | 4M4F / 6M2F / 0-1M6F | 267-279 |
| S7 | 9 m grip strength | 8/16 | 8M8F / 8M8F / 8M8F | 265-287 |
| S8 | 3 and 9 m axonal transport | 6-8 | 5M1F / 6M0F / 3M3F<br>4M4F / 6M2F / 0-1M6F | 91-97<br>267-279 |
| S9 | 9 and 15 m axonal transport | 6-8 | 9 m: n.a. / 6M2F / n.a.<br>15 m: n.a. / 4M2F / n.a. | 267-279<br>450-459 |
| 7 | 9 m sciatic nerve blots | 7-8 | 3M5F / 3M5F / 4M3F | 265-280 |
| 8 | WT axonal transport with TyrRS | 5-8 | PBS: 2M3F / TyrRS<sup>WT</sup>: 0M8F<br>/ TyrRS<sup>E196K</sup>: 0M8F | 58-132 |
| 9A-D | 9 m Hom axonal transport +BDNF | 4 | n.a. / n.a. / 3M1F vs 3M1F | 271-272 |
| 9E-H | 9 m Hom axonal transport +AAV | 4 | n.a. / n.a. / 0M4F vs 0M4F | 273-274 |
| S10 | 9 m +AAV TA blots | 4 | n.a. / n.a. / 0M4F vs 0M4F | 273-274 |
| S11A-D | 9 m Hom axonal transport M vs F | 4-11 | n.a. / n.a. / 4M vs 11F | 267-279 |
| S11E-H | 9 m Hom axonal transport +BDNF | 8-15 | n.a. / n.a. / 4M11F vs 3M5F | 267-279 |

**Table S2.** Primary antibodies used in this study. *BDNF*, brain-derived neurotrophic factor; *ChAT*, choline acetyltransferase; *CST*, Cell Signaling Technology; *DSHB*, Developmental Studies Hybridoma Bank; *DIC*, dynein intermediate chain; *eIF2α*, α subunit of eukaryotic initiation factor 2; *GFP*, green fluorescent protein; *Gt*, goat; *HRP*, horseradish peroxidase; *IF*, immunofluorescence; *Ms*, mouse; *NF*, neurofilament; *Rb*, rabbit; *RILP*, Rab-interacting lysosomal protein; *RRID*, Research Resource Identifier; *Sp*, species; SV, synaptic vesicle; *WB*, western blotting.

| Target | Sp. | Clonality | Company<br>/Reference | Catalogue # | RRID | IF<br>dilution | WB<br>dilution |
| --- | --- | --- | --- | --- | --- | --- | --- |
| BDNF | Rb | Poly | Novus | NB100-98682 | AB_1290643 | n/a | 1:1,000 |
| ChAT | Gt | Poly | Chemicon | AB144P | AB_2079751 | 1:250 | n/a |
| DIC | Ms | Mono | Santa Cruz | sc-13524 | AB_668849 | n/a | 1:500 |
| p-DIC (S81) | Rb | Poly | Mitchell *et al.* [3] | n/a | n/a | n/a | 1:300 |
| p-eIF2α (S51) | Rb | Poly | CST | 9721 | AB_330951 | 1:50 | n/a |
| GFP | Rb | Poly | Abcam | ab290 | AB_303395 | n/a | 1:5,000 |
| Hook1 | Rb | Mono | Abcam | ab150397 | n/a | n/a | 1:1,000 |
| Human IgG Fc<br>(HRP) | Gt | Poly | Abcam | ab98624 | AB_10673832 | n/a | 1:10,000 |
| neurofilament<br>(2H3) | Ms | Mono | DSHB | 2H3 | AB_531793 | 1:250 | n/a |
| NF200 | Ms | Mono | Sigma | N0142 | AB_477257 | 1:500 | n/a |
| peripherin | Rb | Poly | Chemicon | AB1530 | AB_90725 | 1:500 | n/a |
| RILP | Rb | Poly | Abcam | ab140188 | n/a | n/a | 1:1,000 |
| Snapin | Rb | Poly | Granata *et al.* [1] | n/a | n/a | n/a | 1:1,000 |
| SV2 (pan) | Ms | Mono | DSHB | SV2 | AB_2315387 | 1:50 | n/a |
| α-tubulin | Ms | Mono | CST | 3873 | AB_1904178 | n/a | 1:2,000 |
| V5 | Ms | Mouse | Thermo<br>Fisher | R960-25 | AB_2556564 | n/a | 1:5,000 |

**Table S3.** Secondary antibodies used in this study. *Dk*, donkey; *Gt*, goat; *IgG*, immunoglobulin G; *HRP*, horseradish peroxidase; *Ms*, mouse; *Rb*, rabbit; *RRID*, Research Resource Identifier; *Sp*, species.

| Target | Sp. | Conjugate | Company | Catalogue # | RRID | Dilution |
| --- | --- | --- | --- | --- | --- | --- |
| Gt IgG | Dk | AlexaFluor555 | Life Technologies | A-21432 | AB_141788 | 1:500 |
| Ms IgG | Dk | AlexaFluor488 | Life Technologies | A-21202 | AB_141607 | 1:250-500 |
| Ms IgG | Dk | AlexaFluor555 | Life Technologies | A-31570 | AB_2536180 | 1:250-500 |
| Ms IgG | Gt | HRP | Sigma | AP308P | AB_11215796 | 1:5,000 |
| Rb IgG | Dk | AlexaFluor488 | Life Technologies | A-21206 | AB_2535792 | 1:250-500 |
| Rb IgG | Dk | HRP | Bio-Rad | 1706515 | AB_11125142 | 1:5,000 |

## Notes

### Summary of Updates

New data/figures: 2, 5, 8, S1, S9. Updated figure: S12. Updated text and title. New author list.

